# Partitioning Fraction of Variance Explained into Strong Localized Effects and Weak Diffuse Effects

**DOI:** 10.64898/2026.01.06.697735

**Authors:** Fang Nan, David Azriel, Armin Schwartzman

## Abstract

High-dimensional genetic data present substantial challenges for estimating the fraction of variance explained (FVE) by genome-wide single-nucleotide polymorphisms (SNPs). In the context of genetics the VFE is called SNP heritability. Standard approaches for FVE estimation, such as GWAS heritability (GWASH) and linkage disequilibrium score (LDSC) regression, typically assume Gaussian distributions for SNP effect sizes. However, empirical evidence indicates that SNP effects are often heavy-tailed, with a small subset of variants exerting disproportionately large influence. Such settings violate the recently established bounded-kurtosis effect (BKE) condition, under which these FVE estimators are consistent. Consequently, widely used methods may yield severely biased estimates when strong effects are present. We introduce a decomposed FVE estimation framework that accommodates heavy-tailed and heterogeneous SNP effect distributions. The proposed approach partitions total heritability into contributions from strong and weak genetic effects, estimating the former using low-dimensional adjusted *R*^2^ and the latter using an extension of FVE estimation methodology that remains valid under BKE compliance. We further develop a test for detecting violations of the BKE condition and compare several high-dimensional screening procedures for identifying strong-effect SNPs when they are not known in advance. Simulation studies show that the proposed decomposition substantially improves estimation accuracy over existing approaches in the presence of heavy-tailed effects. Application to the Adolescent Brain Cognitive Development (ABCD) Study demonstrates the practical utility of the method, yielding more reliable heritability estimates for the PolyVoxel Score, a neuroimaging-based biomarker linked to iron accumulation. Our results highlight the importance of accommodating effect heterogeneity in large-scale genomic studies.

## 1 Introduction

High-dimensional data, including whole-genome genotyping, exhibits significant complexity, often with dimensionalities that far exceed the number of subjects. As with many complex behavioral traits, individual biomarkers typically contribute only marginally to the overall variance in behavioral phenotypes. However, the vast number of potential biomarkers can collectively make a substantial contribution to the fraction of variance explained (FVE).

Genome-wide Association Studies (GWAS) have become the dominant approach in genetic research on complex disorders due to their relative simplicity, decreasing costs, and notable successes [Visscher et al., 2017]. These studies typically involve regressing quantitative trait measures or case-control status against allele counts for millions of single nucleotide polymorphisms (SNPs) using linear or logistic models. The results are summary statistics—regression coefficients, standard errors, test statistics, and p-values that describe the marginal associations between each SNP and the studied outcome.

A key concept in understanding complex disorders is SNP-heritability, 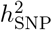, which quantifies FVE attributed to additive relationships among all GWAS SNPs, regardless of their significance [Yang et al., 2010, Lee et al., 2011]. Estimates of 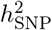 suggest that a considerable portion of susceptibility to complex disorders (approximately 10% - 40%) may be captured by linear additive models based on the same SNPs surveyed by GWAS. The coefficients multiplying the SNP counts in such models and representing the association of each SNP with the outcome are often referred to as SNP effects. However, many existing methods for estimating 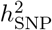 assume a Gaussian distribution for marginal SNP effects, whereas empirical evidence suggests that these effects are not well-characterized by such a distribution [Gusev et al., 2014]. Instead, SNP effects often exhibit a mixture of weak and strong values that do not correspond to a Gaussian distribution [Holland et al., 2020]. This highlights the need for more flexible and realistic methods to model the effect of SNPs. Moreover, current methods often lack robust theoretical foundations and depend on ad hoc simulation studies for validation. Their performance, including bias and error rates, has been shown to be sensitive to the parameters of these simulations [Evans et al., 2018]. To ensure reliable estimates of 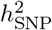, it is crucial to establish realistic conditions and develop methods for testing these conditions with real data.

Estimating the FVE requires advanced statistical methods specifically designed for high-dimensional genetic data. Many FVE estimators exist with various properties, such as consistency [Schwartzman et al., 2019, Dicker, 2014], suitability for non-Gaussian effects [Janson et al., 2017] and non-Gaussian predictors [Hou et al., 2019], applicability to summary statistics [Speed and Balding, 2019, Shi et al., 2016], and effectiveness across both small [Schwartzman et al., 2019, Janson et al., 2017] and large datasets [Speed et al., 2012, Shi et al., 2016]. As a benchmark, an FVE estimator for SNP-based heritability 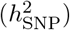 of continuous traits, termed GWAS heritability (GWASH), under a fixed SNP effects framework [Schwartzman et al., 2019], has been developed. However, both GWASH and other summary statistic-based approaches, such as linkage disequilibrium score (LDSC) regression [Bulik-Sullivan et al., 2015], assume that all SNP effects have similar magnitudes. As a result, these methods typically model SNP effects as drawn from a Gaussian distribution centered around zero. As mentioned above, This assumption often does not reflect real-world scenarios [Gusev et al., 2014], where certain SNPs have significantly stronger effects compared to others. This means that the distribution of SNP effects could be heavy-tailed, indicated by a large kurtosis. Although Janson et al. [2017] relaxes the Gaussian-effect assumption and is robust to non-Gaussian signals, their approach requires the predictors to be i.i.d., an assumption rarely satisfied in genetic data where linkage disequilibrium induces correlation among SNPs, thus limiting its practical applicability in this context.

A high kurtosis of SNP effects violates an important condition, shown by Azriel et al. [2025] to be necessary and sufficient for consistent FVE estimation using GWASH and LDSC regression. Azriel et al. [2025] formally defined this condition, referred to as bounded-kurtosis effects (BKE), requiring the kurtosis of the distribution of SNP effects to be asymptotically bounded as the number of predictors, i.e., SNPs, increases. In this article, we first extend the GWASH approach by developing a more general framework for FVE estimation that accommodates heavier-tailed, non-Gaussian distributions of SNP effects. Violations of the BKE condition often occur when a small subset of SNPs exerts disproportionately large effects, giving rise to a mixture structure in which strong and weak effects coexist. By explicitly modeling this structure, we account for scenarios where SNP effects are partially concentrated in a small subset of variants while remaining broadly distributed across the genome.

To address the violation of the BKE condition, we propose a decomposed FVE estimator. The essential idea is to quantify the relative contributions of different groups of predictors, a concept commonly referred to as enrichment in the context of GWAS [Finucane et al., 2015]. This approach partitions the total FVE into contributions from strong and weak genetic effects, allowing their heritability to be estimated in sequence and then assembled to obtain the total SNP heritability. After decomposition, we assume that the BKE condition holds within the high-dimensional weak component. The strong-effect component is low-dimensional and therefore does not require the BKE condition. This procedure is conceptually analogous to ANOVA in linear regression, but is adapted here to handle high-dimensional genetic predictors.

In our framework, the number of SNPs with strong effects is assumed to be smaller than the number of subjects, resulting in a low-dimensional setting. Consequently, the heritability attributable to strong effects can be reliably estimated using the adjusted *R*^2^. In contrast, the number of weak-effect SNPs typically exceeds the sample size, creating a high-dimensional estimation problem. For this component, we apply the GWASH method [Schwartzman et al., 2019] and Genome-wide Complex Trait Analysis (GCTA, Yang et al. [2011]) for comparison. LDSC regression with fixed intercept gives values that are numerically very close to GWASH [Schwartzman et al., 2019, Azriel et al., 2025], and we also included it in our analysis. Although GCTA is one of the most widely used methods for estimating SNP heritability, its robustness to heavy-tailed SNP effect distributions has not been rigorously examined in theory. Our simulation studies, however, indicate that GCTA is also sensitive to violations of the BKE condition. Encouragingly, integrating GCTA within our proposed decomposition framework substantially improves its performance under non-Gaussian settings, enhancing estimation accuracy even when the BKE condition is violated. By modeling the strong and weak effects separately and then aggregating their contributions, our method effectively mitigates the impact of BKE violations and yields a consistent estimate of the total SNP heritability.

Sometimes, SNPs with strong effects are already known from previous studies. However, when such a subset is not known a priori, identifying these strong effects becomes a crucial step in our framework. We first develop a testing procedure to evaluate whether the SNP effects in the sample satisfy the BKE condition, thereby informing the suitability of applying the proposed decomposed FVE estimation. Next, we propose to use high-dimensional screening methods for this purpose. Existing screening methods for high-dimensional data have not been implemented for FVE decomposition before, and they were originally designed for sparse models with many zero coefficients. For instance, the Sure Independence Screening (SIS, [Fan and Lv, 2008]) procedure screens out irrelevant features based on their marginal correlations with the response variable. The Robust Rank Correlation Screening (RRCS, [Li et al., 2012]) method employs Kendall’s *τ* rank correlation to select variables in ultra-high-dimensional data, offering robustness to outliers and invariance to monotonic transformations. The High-dimensional Ordinary Least Squares Projection (HOLP, [Wang and Leng, 2016]) method ensures consistency in variable selection without relying on strong assumptions like those in SIS. Additionally, the Bonferroni correction [Bonferroni, 1936] can be readily applied in our setting to identify SNPs with strong effects. We present a simulation study to evaluate and compare the performance of the four aforementioned methods in connection to FVE estimation. A repository containing the code used to run the simulation analyses in this section is available at: https://github.com/Fangn06/Decomposed-FVE-Estiamtion.

To demonstrate the practical utility of our proposed framework, we apply it to the Adolescent Brain Cognitive Development (ABCD) Study [Casey et al., 2018], one of the largest longitudinal studies of brain development and child health in the United States. The ABCD dataset includes extensive whole-genome genotyping and rich behavioral and cognitive phenotypes from over 11,000 children who were 9–10 years old at baseline, with their biological and behavioral development longitudinally tracked over subsequent years [Luciana et al., 2018, Casey et al., 2018]. Genetic factors significantly influence mental illnesses [Laville et al., 2019, Polderman et al., 2015], and understanding these genetic contributions is crucial for elucidating the etiology of psychiatric disorders and developing targeted treatments [Sullivan and Geschwind, 2019]. The primary outcome in our analysis is the PolyVoxel Score (PVS), defined by Loughnan et al. [2024], which measures how closely an individual’s brain pattern matches the archetypal hemochromatosis brain, marked by regional iron accumulation in motor circuits. In the ABCD dataset, our BKE condition test flagged potential violations in specific chromosomes, pointing to the presence of strong SNP effects that contribute to the heritability of the PVS. Notably, the decomposed estimator yields larger FVE estimates than its non-decomposed counterpart across the majority of chromosomes, suggesting that explicitly accounting for strong genetic effects affords a more complete characterization of the genetic architecture underlying the PVS phenotype.

## 2 General Setting

### 2.1 Polygenic Model and FVE

Suppose a continuous outcome *y*_*i*_ is measured with a panel of *m* predictors 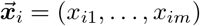 in *n* independent and identically distributed subjects, *i* = 1, …, *n*. In the context of genetics, these are SNP allele counts (taking values 0, 1, 2). The poly-additive model, called polygenic model in genetics ([Fisher, 1919, Lynch et al., 1998]), in scalar or vector form, is

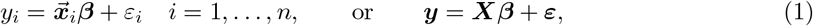

where ***y*** = (*y*_1_, …, *y*_*n*_)^⊤^, ***β*** = (*β*_1_, …, *β*_*m*_)^⊤^, ***X*** is the regression matrix with rows 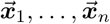 representing subjects’ SNP panels and columns ***x***_1_, …, ***x***_*m*_ representing SNPs for all subjects. The errors ***ε*** = (*ε*_1_, …, *ε*_*n*_)^⊤^ are independent of ***X***, and *ε*_*i*_ are IID with zero mean and variance 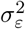. To simplify the notation and theoretical calculations, we assume 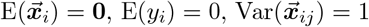 for *j* = 1 …, *m*, and Var(*y*_*i*_) = 1. Let 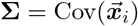 denote the *m* × *m* covariance matrix among SNP predictors in the underlying population.

The goal is to estimate the SNP heritability or FVE of Model (1). When Model (1) is treated as a fixed-effects model, where the SNP effects ***β*** are considered fixed, the FVE, denoted by 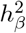, is defined as

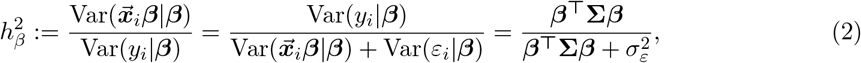

also referred to as the “fixed-effects heritability”. Here, we define the conditional variance of *y*_*i*_ as 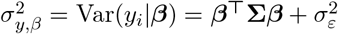.

However, it is often more mathematically convenient to treat Model (1) as a random-effects model, where the SNP effects ***β*** are considered random. In this case, the FVE is referred to as the “random-effects heritability”, denoted by *h*^2^. Here we assume that the SNP effects *β*_1_, …, *β*_*m*_ are i.i.d. with mean zero and a common variance Var(*β*_*j*_) = *h*^2^*/m*. We also assume that ***β*** is independent of both 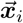 and *ε*_*i*_. Under these assumptions, we have E(***β***^⊤^**Σ*β***) = *h*^2^ and Var(*ε*_*i*_) = 1 − *h*^2^. Therefore, the quantity *h*^2^ represents the unconditional FVE under Model (1):

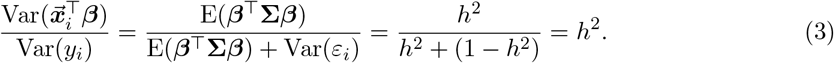

To model the SNP effects ***β*** as non-Gaussian, an important concept in this context is kurtosis. Treating the SNP effects as random, the kurtosis of *β*_*j*_ is defined as

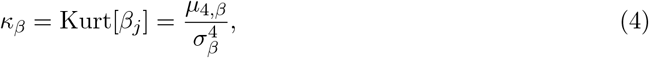

where *µ*_4,*β*_ is the fourth central moment and *σ*_*β*_ is the standard deviation of *β*_*j*_. Typically, the quantity Kurt[*β*_*j*_] − 3 is referred to as the excess kurtosis of *β*_*j*_, using the normal distribution, whose kurtosis is 3, as the reference.

### 2.2 GWASH Estimator

Considering Model (1) and define ***η*** = (*η*_1_, …, *η*_*m*_) as the vector of correlations between the entries of 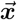 and *y* given the SNP effects. Specifically, ***η*** is given by:

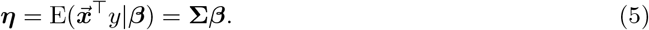

Let *s*^2^ = ||***η***||^2^ and *µ*_2_ represent the second spectral moment of the correlation matrix **Σ** of 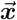, defined as *µ*_2_ = tr(**Σ**^2^)*/m*. Using these definitions, we have that

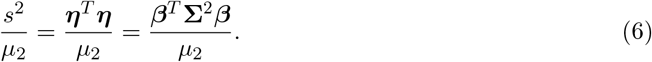

Recall that *β*_1_, …, *β*_*m*_ are i.i.d. with mean zero and variance Var(*β*_*i*_) = *h*^2^*/m*, hence,

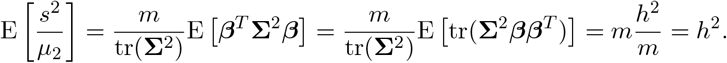

When considering standardization in the sample, the predictors ***X*** and the outcome ***y*** are transformed into their standardized versions as follows:

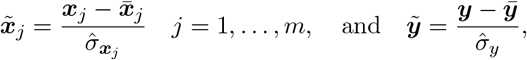

where 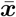 and 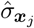 are the sample mean and standard deviation of 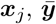 and 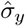 are the sample mean and standard deviation of the outcome ***y***. Let 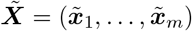 represent the standardized version of ***X***, obtained by standardizing its columns.

The sample correlation between ***x***_*j*_ and ***y*** is then computed as:

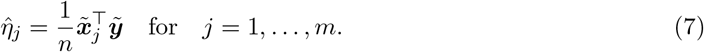

which serves as the sample version of ***η*** defined in (5). The *j*-th LD-score, for *j* = 1, …, *m*, as defined by [Bulik-Sullivan et al., 2015], measures the sum of squared sample correlations between ***x***_*j*_ and all other predictors. The population and sample versions of the LD score are defined, respectively, as

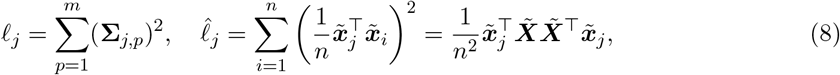

Notice that since we assume Var(*X*_*ij*_) = 1, it follows that **Σ** is the population correlation matrix of the predictors. The bias-corrected LD-score 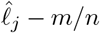 is approximately unbiased for *ℓ*_*j*_ for Gaussian predictors as shown in Equation (5) of [Azriel et al., 2025].

The GWASH estimator can be obtained as a plug-in estimator of Equation (6):

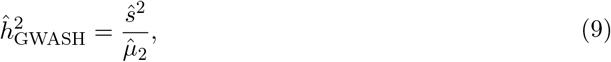

where 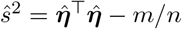 is approximately an unbiased estimator of *s*^2^ as shown in Equation (6) of [Azriel et al., 2025], and 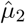 is an unbiased estimator of the second spectral moment 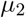:

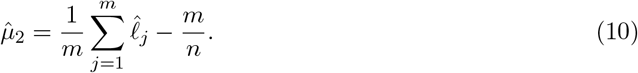

## 3 Estimation Consistency and the Effect of Kurtosis

### 3.1 Consistency Conditions of GWASH Estimator

Two important conditions for the consistency of the GWASH estimator and LDSC regression with fixed intercept were discussed by [Azriel et al., 2025]. The first is the weak dependence condition, which requires that the second spectral moment of the LD matrix **Σ** to be bounded. This condition is generally assumed to hold in GWAS as dependence between SNPs is local [Xie et al., 2011]. However, it may be violated in heterogeneous populations, in which case the issue can be mitigated by first regressing the outcome on principal components that capture population structure [Singh Sachan, 2025]. The focus of the current paper is on the second consistency condition, namely, the bounded-kurtosis-of-effects (BKE) condition, which is formally defined as:

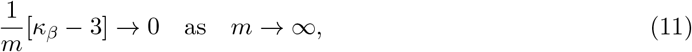

where *κ*_*β*_ is the kurtosis of *β*_*j*_ as defined in Equation (4).

While a detailed description of the BKE condition can be found in [Azriel et al., 2025], here we briefly discuss the implications of that in our setting. Let

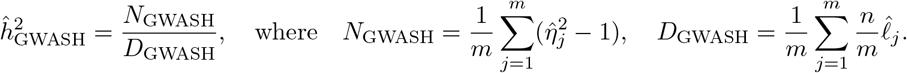

Under Model (1), and assuming weak dependence together with additional regularity conditions (e.g., bounded moments of the error term *ε*) as specified in Equation (12) of Azriel et al. [2025], we have

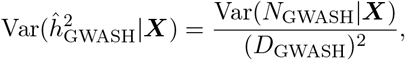

where

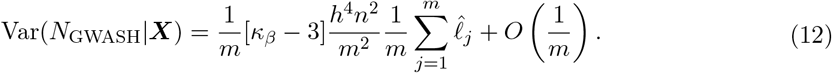

The expression above in (12) explains how the BKE condition guarantees the consistency of 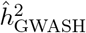. Here, we assume that *m/n* converges to a positive constant as both *m* and *n* increase, while the average of 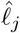’s converges to a constant under the assumption of weak dependence. The expression in the square brackets is the excess kurtosis. If ***β*** follows a normal distribution, this excess kurtosis is zero, meaning that the term does not contribute to the variance and the variance goes to zero at a rate of 1*/m* given by the residual term. However, if ***β*** is not normally distributed, the excess kurtosis may be large, and the variance may not go to zero. We show an example in the next section.

### 3.2 Kurtosis and Modeling of Coefficient Effects

To more thoroughly investigate and model SNP effect coefficients ***β*** with varying kurtosis, we present here a Gaussian mixture model as an example, which is implemented in our subsequent simulation studies. However, we note that this is just one possible method for modeling the effects, and alternative approaches may also be considered.

Consider a Gaussian mixture distribution of ***β***, where a small fraction *p* = *κ/m* of *β*’s having variance *ψ*^2^ and the rest have variance *γ*^2^, and *p* ∈ [0, 1].

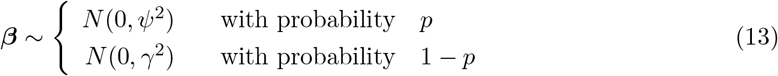

The variance of *β* is 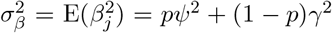. By letting the total variance of 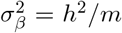, we have 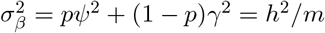, with *pψ*^2^ = *θh*^2^*/m* and (1 − *p*)*γ*^2^ = (1 − *θ*)*h*^2^*/m*, where *θ* denotes the fraction of variance that the small fraction *β*’s take. Then with restricting *p* ∈ (0, 1), we have *ψ*^2^ = *θh*^2^*/*(*mp*) and *γ*^2^ = (1 − *θ*)*h*^2^*/*[*m*(1 − *p*)]. In order to derive the kurtosis in terms of *p* and *θ*, we first derive the fourth moment of *β* as

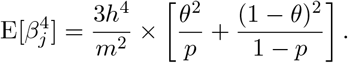

Then, since 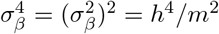, the kurtosis is

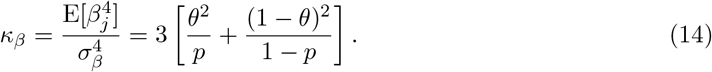

The mixture Gaussian model for ***β*** provides a flexible framework that accommodates both scenarios where the BKE condition is satisfied and those where it is violated. As shown in Equation (14), when *p* = *θ*, the mixture model reduces to a standard Gaussian distribution, which satisfies the BKE condition. Furthermore, each component in the mixture individually satisfies the BKE condition when considered in isolation. However, by manipulating the value of *p* and *θ*, for example, setting *p* to a relatively small value and *θ* to a large proportion, the resulting distribution exhibits high kurtosis, thereby violating the BKE condition, as demonstrated by the following proposition. The corresponding proof of this proposition is provided in Appendix B.1.

#### Proposition 3.1.

*Under the mixture Gaussian model in (13), suppose θ* = *θ*_0_ ∈ (0, 1) *is fixed and p* ∈ [0, 1]. *Then the BKE condition is violated if either mp* → *c <* ∞ *or m*(1 − *p*) → *c <* ∞ *as m* → ∞. *The BKE condition is satisfied when p* = *θ*_0_, *or when p* ∈ {0, 1}, *or when θ*_0_ ∈ {0, 1}, *in which case the distribution reduces to a standard Gaussian*.

### 3.3 Relationship Between the Kurtosis of *β* and *η*

In practice, the true coefficients ***β*** are unobservable; therefore, the exact value of *κ*_*β*_ is not directly available. In such cases, it is important to understand the relationship between *κ*_*β*_ and 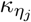, the latter being estimable from observable data. Here, 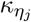 denotes the kurtosis of *η*_*j*_, defined as

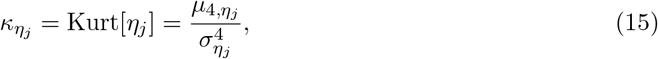

where 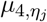 is the fourth central moment and 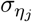 is the standard deviation of *η*_*j*_. This relationship between *κ*_*β*_ and 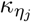 is established by the following proposition:

#### Proposition 3.2.

*Consider Model (1) and assume the random effects* 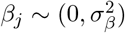. *Then*

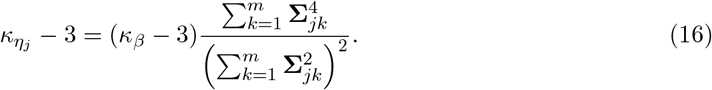

Proposition 3.2 shows that although the kurtosis of *β*_*j*_ is not equal to the kurtosis of *η*_*j*_, a higher value of 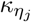 generally indicates a higher value of *κ*_*β*_. The proof of this proposition is provided in Appendix B.2. The effect of the dependence in **Σ** on this relationship is demonstrated through simulations in Section 6.2.

## 4 FVE Decomposition

If the BKE condition is violated, the GWASH estimator and others become inappropriate due to its inconsistency. To address this, we propose an alternative method for estimating FVE, referred to as FVE decomposition, and subsequently introduce the decomposed FVE estimator.

The core idea is to determine the relative contribution of different groups of predictors, by partitioning the total FVE among subsets of predictors. In our framework, SNP effects are partitioned into two components: a low-dimensional subset and a high-dimensional subset, with the latter assumed to satisfy the BKE condition. The FVE attributable to the low-dimensional component can be accurately estimated using the adjusted *R*^2^. For the high-dimensional component, we employ high-dimensional FVE estimators, such as GWASH or GCTA, to quantify its contribution to the total FVE, as the BKE condition is presumed to hold after partitioning. A brief description of GCTA is presented in Appendix A.1.

### 4.1 FVE Decomposition into Sub-Classes of Predictors

Here, we propose a general FVE decomposition framework that partitions the total FVE into two components of arbitrary dimensionality. In this section, we describe the decomposition at the population level using theoretical parameters. The estimation procedures corresponding to this decomposition will be detailed in the next section.

In model (1), consider a random row vector 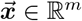 and a random variable *y* ∈ ℝ, both with mean zero, dropping the subject index *i* for simplicity. The vector of effects ***β*** can be characterized as providing the best linear approximation of *y* by 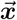:

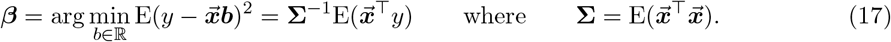

Consider a partition of the predictors 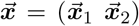 into two subsets, where 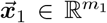 and 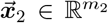, partitioning the effects vector as 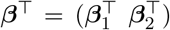 and the covariance **Σ** into blocks 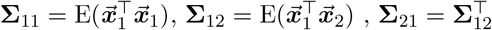 and 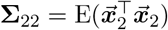. Based on the above partition, we can derive the decomposition of FVE as follows.

#### Proposition 4.1.

*Consider Model (1) and let* 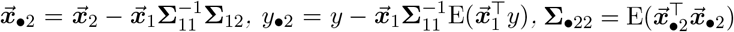. *Then the conditional FVE in (2) can be decomposed as*

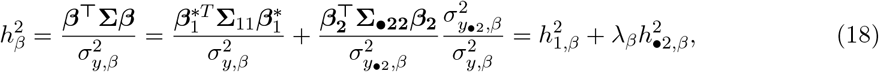

*where* 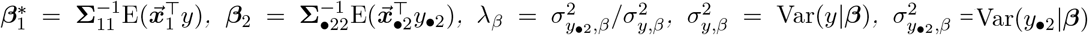

The first term in the sum (18) relates to the marginal regression of *y* on 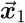, while the second term corresponds to the regression of *y* on 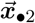, adjusted for 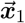. Notice that the two terms correspond to orthogonal predictors so that they can be estimated separately and added. The detailed proof of this proposition is provided in Appendix B.3.

### 4.2 Estimation of FVE Components

In this section, we provide the estimation details for Proposition 4.1. Let 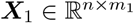 and 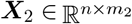 denote the partitioned SNP matrices, where *n* is the number of subjects and *m*_1_+*m*_2_ = *m > n* is the total number of SNPs. Let ***y*** ∈ ℝ^*n*^ be the trait of interest. We assume that *m*_1_ *< n*, so the estimation of 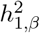, as defined in equation (18), corresponds to a low-dimensional setting. Therefore, it can be reliably estimated using the adjusted *R*^2^, which we denote by 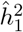,

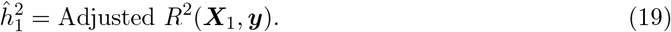

To isolate the contribution of the high-dimensional component ***X***_2_, we residualize both ***X***_2_ and ***y*** with respect to ***X***_1_. Specifically, we compute the residualized matrices:

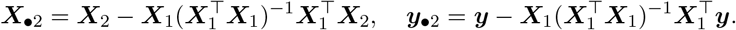

Note that 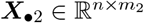, and since *m*_2_ *> n*, estimating 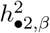 is a high-dimensional problem. This can be addressed using high-dimensional FVE estimation methods, such as GWASH, LDSC, or GCTA. In this case, the estimate of the residual heritability is denoted by 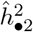,

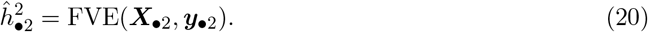

Moreover, the ratio *λ*_*β*_ in equation (18) can be easily estimated by the sample variance ratio 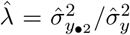, where 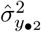 and 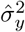 denote the sample variances of ***y***_•2_ and ***y***, respectively. Finally, by substituting the estimates 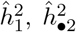, and 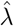 into equation (18), we obtain the estimated FVE for all SNPs 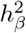, denoted by *ĥ*^2^. A pseudo-algorithm outlining the estimation procedure for FVE decomposition is provided in Algorithm 1.

#### Algorithm 1

FVE Decomposition Estimation Procedure

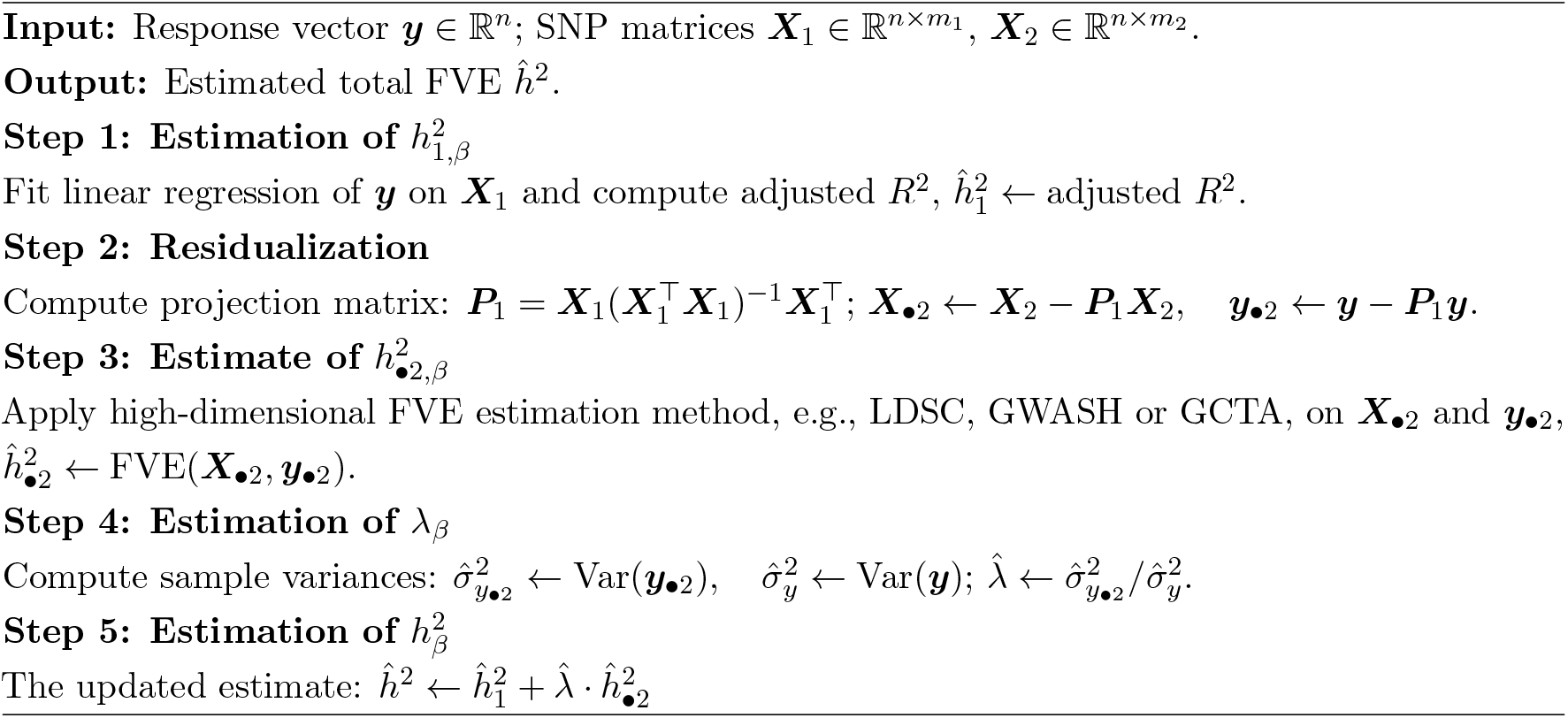

## 5 Testing the BKE Condition on Effect Coefficients

We consider now Model (1) and assume that *β*_1_, …, *β*_*m*_ are IID with mean zero and variance of *h*^2^*/m*. To more effectively evaluate whether the BKE condition holds, we formulate the following hypothesis test for the effects *β*_*j*_’s:

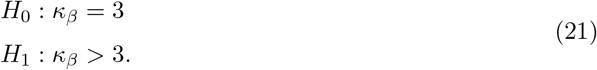

In practice, this hypothesis is not directly testable because the true effects vector ***β*** is not observable. To address this limitation, we instead base the following proposition on the estimable correlation vector ***η***, as introduced in Equation (7), to test the hypothesis in terms of *κ*_*β*_ as stated in (21).

### Proposition 5.1.

*Consider Model (1) and assume the random effects* ***β*** ~ *N*(0, *h*^2^***I****/m*). *Then the covariance matrix of the sample correlation vector* 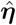 *given* ***X*** *is*

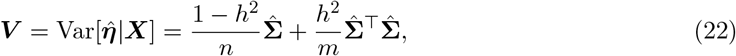

*where* 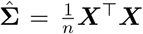 *is the sample covariance matrix of* ***X***. *Furthermore, assuming errors* 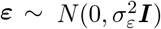 *the sample correlation vector* 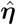 *follows a multivariate normal distribution:*

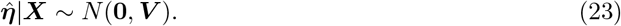

The proof of this proposition can be found in Appendix B.4. This proposition facilitates testing the null hypothesis that the BKE condition holds. To construct a test statistic, we whiten 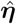 using the spectral decomposition of 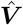, where 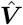 is the same as ***V*** but with *ĥ*^2^ replacing *h*^2^. Let 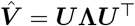 be the eigendecomposition, and let *r* be the rank of 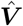. Define the whitened correlation as

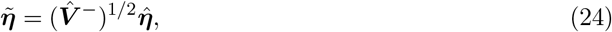

where 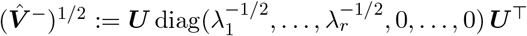 is the Moore-Penrose square-root inverse. Under the BKE condition, the effective components 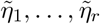 are i.i.d. standard normal.

We then apply the Jarque-Bera test to the *r* effective whitened scores 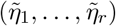, which jointly evaluate whether their sample skewness and excess kurtosis are consistent with those of the standard normal distribution. Combined with the results in Section 3.3, a large kurtosis in 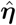 generally implies elevated kurtosis in ***β***. Therefore, rejection of the Jarque–Bera null provides indirect evidence of a departure from the BKE condition, i.e., *κ*_*β*_ *>* 3.

## 6 Numerical Studies

In this section, we conduct numerical experiments to evaluate our proposed framework, including decomposed FVE estimation, testing for the BKE condition, and screening methods. The structure of this section is as follows. Section 6.1 introduces a numerical approach to control the kurtosis of ***β***, which we then use in all the consequent simulations. In Section 6.2, we examine the relationship between *κ*_*β*_ and 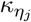, and empirically quantify the findings described in Proposition 3.2. Section 6.3 evaluates the performance of the decomposed FVE estimator. Section 6.4 assesses the performance of the BKE testing procedure. Finally, Section 6.5 investigates the effectiveness of several existing screening methods (detailed in Appendix A.2) and provides a comparative analysis of their performance. A repository containing the code used to run the simulation analyses in this section is available at: https://github.com/Fangn06/Decomposed-FVE-Estiamtion.

### 6.1 Controlling Kurtosis of Effects in Simulation Studies

Due to sampling variability, directly simulating ***β*** from the Gaussian mixture model in Equation (13) makes it difficult to achieve the target kurtosis. To address this issue, we propose a data generation mechanism that ensures the empirical kurtosis of the simulated ***β*** aligns with the specified value in the simulation design.

The basic idea of the mechanism is that we decompose ***β*** into two components (***β***_1_, ***β***_2_) as described in Section 3.2. The strong effects 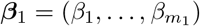 having a common variance *ψ*^2^, and the weak effects 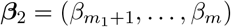 having a common variance *γ*^2^. Then we can find appropriate *ψ*^2^ and *γ*^2^ to ensure the ***β*** has the desired kurtosis. Based on the derivation of *κ*_*β*_ in (14), we can first obtain *ψ*^2^ = (1 − *θ*)*h*^2^*/*[*m*(1 − *p*)] with the given *θ, h*^2^ and *p*. Then we generate ***β***_1_ and ***β***_2_ from the standard normal distribution and then standardize them to ensure these two components have a mean of 0 and variance of 1, respectively. We assume the true Kurt[***β***] is *K* and the classic formula of kurtosis:

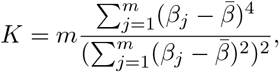

where 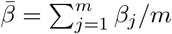. Since ***β***_1_ and ***β***_2_ have been centered and scaled, we can solve the following equations to approximate *γ*^2^:

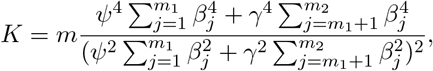

where *m*_1_ + *m*_2_ = *m*. This equation is a quadratic equation and can be solved either with numeric optimization or the quadratic formula 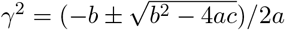, where 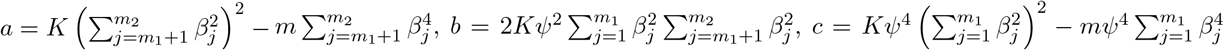. Here we should only consider the positive solution of *γ*^2^. Then the generated ***β*** with specific kurtosis can be expressed as ***β*** = (*ψ****β***_1_, *γ****β***_2_).

### 6.2 Simulation Study I: Relationship Between Kurt[*β*] and Kurt[*η*]

To demonstrate the relationship between Kurt[***β***] and Kurt[***η***], we conducted a series of simulations. We varied the AR(1) correlation parameter *ρ* of **Σ** across the values *ρ* = 0, 0.5, 0.9. The kurtosis of ***β*** were set to 3, 5, 20, 50, and 100, controlled using the method described in Section 6.1. We set *m* = 400, *n* = 200, *θ* = 0.9, *h*^2^ = 0.5, 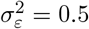. The indices of strong effects are assigned to be from *j* = 201 to *j* = 210.

In this simulation, for each specified kurtosis level of ***β***, we first compute the theoretical kurtosis of ***η*** using Equation (16), which is depicted as the solid line in Figure 1. We then generate ***β*** with the target kurtosis using the method described in Section 6.1, compute ***η*** = **Σ*β*** based on the given **Σ** and ***β***, and compute the sample kurtosis of ***η***. This sample kurtosis is treated as the empirical kurtosis of ***η***, shown as the dashed line in Figure 1.

**Figure 1:**
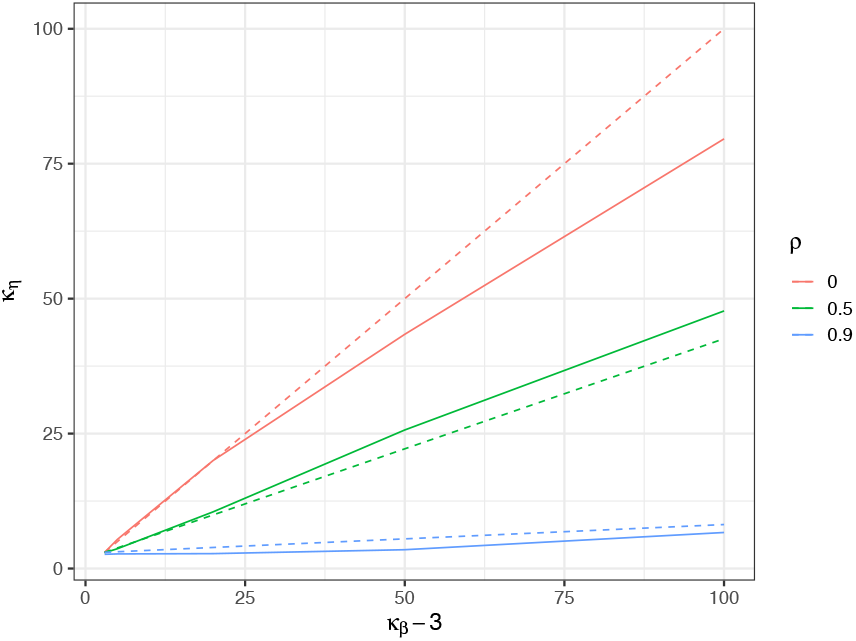
Relationship between *κ*_*β*_ and 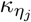. The solid line depicts the theoretical kurtosis of ***η*** as a function of the kurtosis of ***β***, computed from Equation (16). The dashed line shows the empirical kurtosis of ***η*** based on simulated ***β*** with the corresponding kurtosis.

It is worth noting that the theoretical ratio exhibits a boundary effect for *ρ* = 0.5 and 0.9 when the index *j* is close to 0 or *m*. In contrast, the ratio remains stable for interior values of *j*. To avoid this boundary effect, we compute the theoretical kurtosis of *η*_*j*_ at *j* = *m/*2 using Equation (16). For the empirical vector ***η***, we extract a subset from *j* = 50 to *j* = 350 and use this subset to calculate its empirical kurtosis.

The final values of 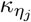 were averaged over 500 simulation iterationsn for each given *κ*_*β*_. The results are presented in Figure 1, where the dashed line represents the theoretical 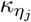 derived from Proposition 3.1, and the solid lines show the empirically averaged 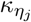 from the simulation. We can observe that for different *ρ*, the empirical results match the trend of the theoretical predictions. These results illustrate the relationship between *κ*_*β*_ and 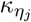 under varying correlation levels. When *ρ* = 0 or 0.5, 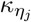 closely tracks *κ*_*β*_, with larger values of 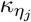 consistently corresponding to larger values of *κ*_*β*_. For *ρ* = 0.9, although the increasing trend becomes less pronounced, the two quantities still exhibit a similar overall pattern.

### 6.3 Simulation study II: Decomposed FVE estimator

Since the overall framework of the proposed decomposed FVE estimation is not limited to high-dimensional settings, we provide results in both low and high-dimensional settings and evaluate their performance accordingly. In the low-dimension setting, we vary *m* = 50, 100, 150 and fix the ratio between *m* and *n* as *n* = 5*m*. We change the kurtosis such that *κ*_*β*_ = 3, 5, 10, 20, 30 for each *m*. The strong effects are the first 5 coefficients in this low-dimensional setting, resulting in *p* = 5*/m*, and we set *θ* = 0.9 and *h*^2^ = 0.5. In the high-dimension setting, we vary *m* = 400, 800, 1200 and fix the ratio between *m* and *n* as *n* = *m/*2. We change the kurtosis such that *κ*_*β*_ = 3, 5, 20, 50, 100 for each *m*. The strong effects are the first 10 coefficients in this high-dimensional setting, resulting in *p* = 10*/m*, and we keep *θ* = 0.9 and *h*^2^ = 0.5. For both scenarios, the distribution of ***X*** is assumed to follow a multivariate Gaussian with the correlation structure being AR(1) (*ρ* = 0, 0.5, 0.9). The number of simulation instances is 1000 in the following studies.

Due to sampling variability, the realized FVE, denoted as 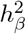 in Equation (2), may deviate from the target FVE *h*^2^ specified in the simulation setup. Moreover, fluctuations in the realized FVE may occur across repeated simulation iterations. Although Section 6.1 presents a method for controlling the kurtosis of ***β***, we find that it is not possible to simultaneously control both the kurtosis of ***β*** and the realized FVE. To quantify the accuracy and stability of the simulations, we use the relative root mean square error (RMSE) as a performance metric. As in the Equation (2), let

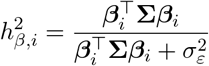

represent the realized FVE conditional on ***β***_*i*_ in the *i*-th simulation realization, and 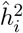 denote the estimated FVE in the same realization, for *i* = 1, …, *M*, where *M* denotes the total number of simulation realizations. The relative RMSE of the estimators is then calculated as:

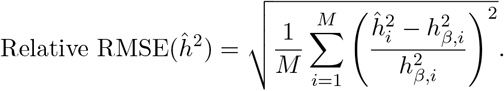

Similarly, we quantify the relative bias as

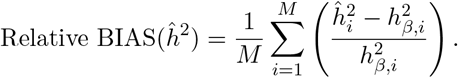

To implement our FVE decomposition, we estimated the explained variance in the low-dimensional component, 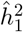 in Equation (19), using the adjusted R^2^, while the remaining component, 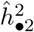 in Equation (20), was estimated using high-dimensional FVE estimators, such as GWASH, GCTA, or LDSC.

The estimation results of the decomposed GWASH estimator relative to the original GWASH estimator are presented in Figure 2 in terms of relative RMSE, and in Figure 11 in Appendix C in terms of relative bias. For both low- and high-dimensional settings, we observe that larger values of *m* lead to lower relative RMSE, while increasing kurtosis in ***β*** results in higher RMSE. Across all settings, the decomposed estimator consistently achieves lower RMSE than the original GWASH estimator, with the improvement being more pronounced at larger values of *ρ*. These results highlight the decomposed estimator’s superior ability to mitigate the adverse effects of large kurtosis, which can otherwise violate the BKE assumptions underlying the original GWASH approach.

**Figure 2:**
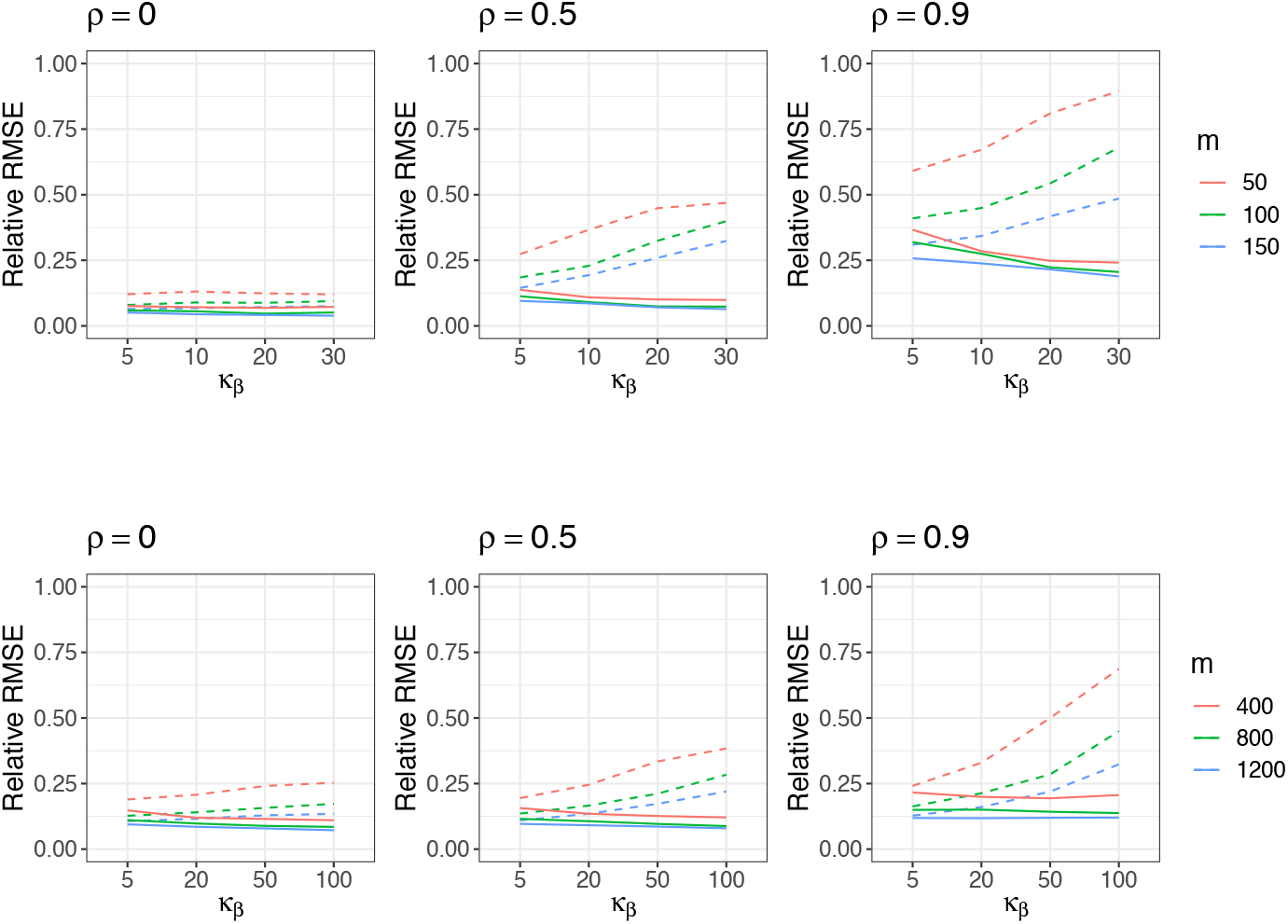
Estimation performance of GWASH without decomposition (dashed line) and decomposed GWASH (solid line) in the low-dimensional setting (top row) and high-dimensional setting (bottom row). The standard error from 1000 simulation instances is about 0.0003 (low-dimension) and 0.0007 (high-dimension). Note that the kurtosis *κ*_*β*_ is configured differently under low-versus high-dimensional settings and hence the scales of the *x*-axis in the top and bottom rows differ.

The results of LDSC and GCTA are similar to those of GWASH, indicating that the decomposed estimator outperforms the original estimator without decomposition under high-kurtosis settings. Further details of the LDSC and GCTA results are provided in Appendix C.

### 6.4 Simulation Study III: Testing the BKE Condition

We evaluated the performance of the proposed test as formulated in Equation (21) and described in Section 5, under both low- and high-dimensional settings. The results are presented in Figure 3. In both cases, the type I error rate (under *κ*_*β*_ = 3) remains close to the nominal level of 0.05 across different values of *m* and *ρ*. The power increases with higher kurtosis, and larger values of *m* generally lead to greater power. Notably, under weak correlation (*ρ* ∈ {0, 0.5}), the test’s power approaches 1 when *κ*_*β*_ reaches 20 in the low-dimensional setting and 100 in the high-dimensional setting.

**Figure 3:**
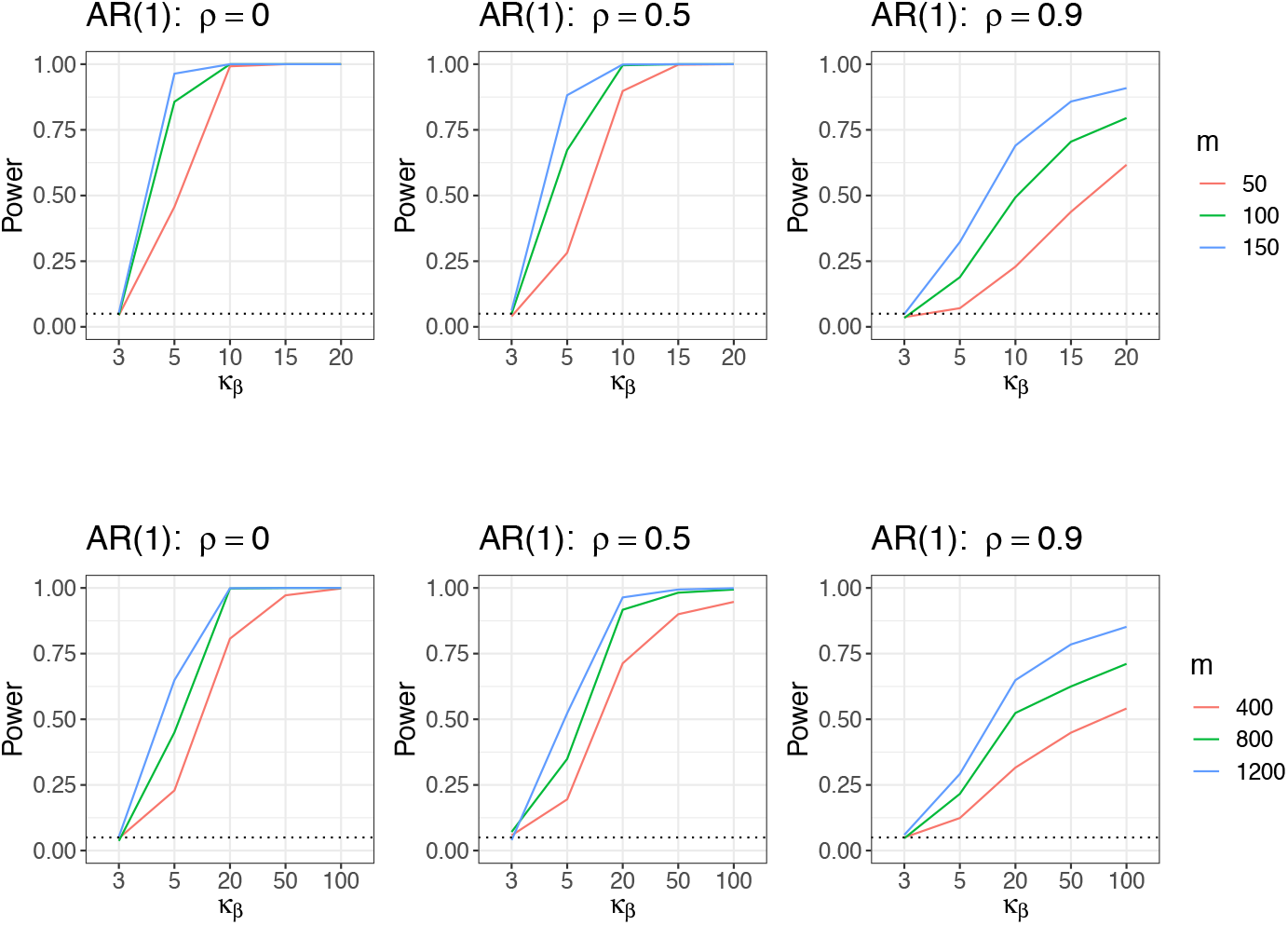
Performance of testing of the BKE condition in the low-dimensional setting (top row) and high-dimensional setting (bottom row). The Type-I error is the value of the power at *κ*_*β*_ = 3. Note that the kurtosis *κ*_*β*_ is configured differently under low-versus high-dimensional settings and hence the scales of the *x*-axis in the top and bottom rows differ.

### 6.5 Simulation study IV: Screening Methods for Identifying Strong Localized Effects

We now study different screening methods through simulations. We fix *m* = 400 and *n* = *m/*2 = 200. The correlation matrix **Σ** follows an AR(1) structure with *ρ* ∈ {0, 0.5, 0.9}. We varied the kurtosis to obtain *κ*_*β*_ ∈ {5, 20, 50, 100} and evaluated five screening methods described in Appendix A.2. As benchmarks, we included two additional approaches: no screening, and an oracle method that leverages the true indices of strong effects to perform the decomposition. To obtain unbiased estimates and to avoid overfitting, the data is split into two parts: 15% of the original sample is used as the screening set to identify strong effects, and the remaining 85% is reserved as the estimation set.

The results of the estimation using different screening methods are presented in Figure 4. For all estimation methods (GWASH, LDSC, and GCTA), the estimation without any screening shows the largest RMSE, while the oracle has the smallest RMSE, reflecting its superior estimation performance due to the use of true indices for strong effects. For GWASH and LDSC, when *κ*_*β*_ is relatively small, all screening methods perform similarly. As *κ*_*β*_ increases, some divergence among the screening methods emerges, although the differences remain modest. For GCTA, the differences in RMSE across the various methods are minor. Summarizing the performance of all screening methods across GWASH, LDSC, and GCTA, we observe that HOLP consistently achieves the best performance and is therefore recommended.

**Figure 4:**
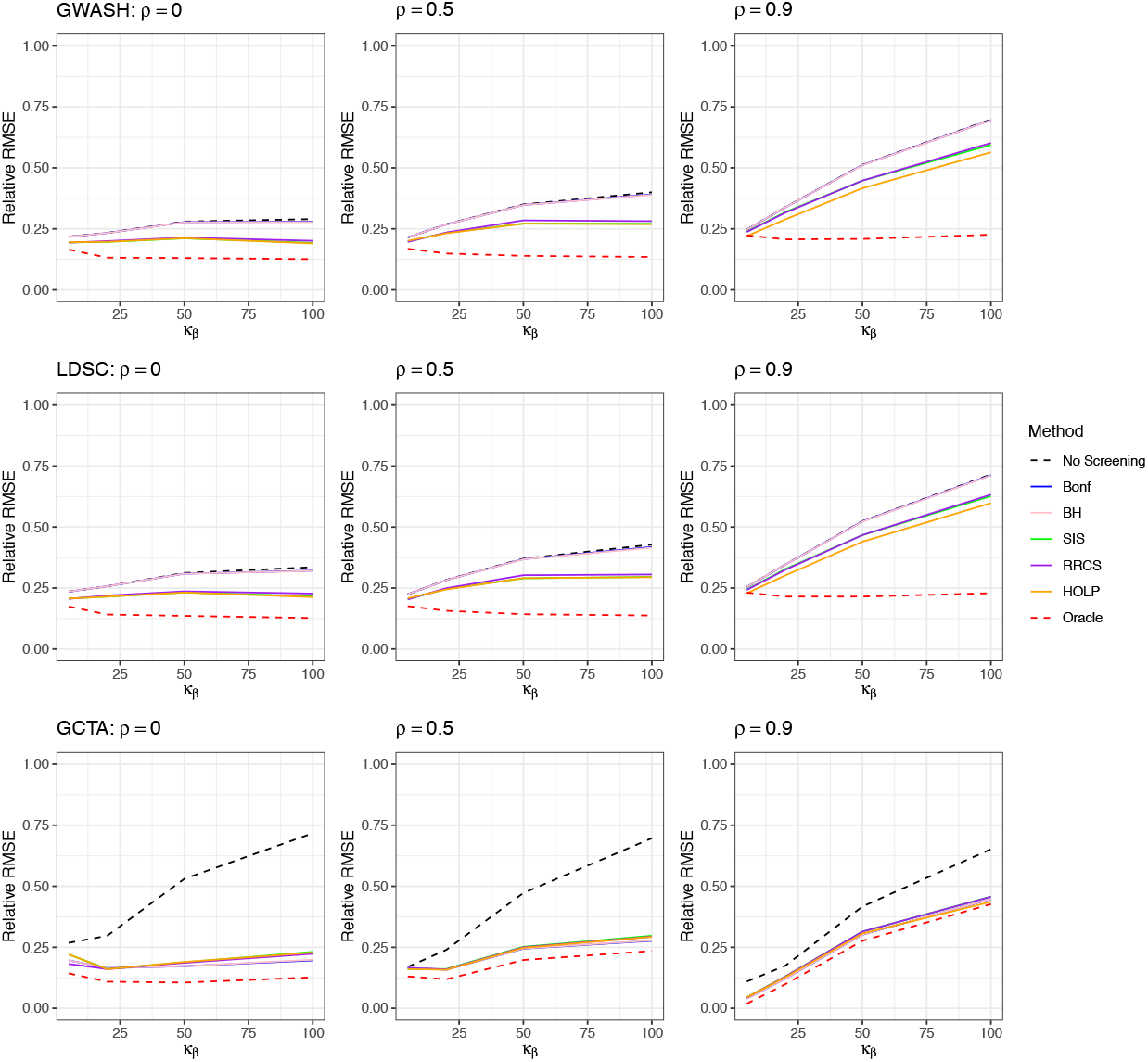
Estimation performance across different screening methods. Results for GWASH, GCTA, and LDSC are displayed in the panel from the top to the bottom, respectively. The standard error from the 1000 simulations instances are about 0.001 (GWASH), 0.0007 (GCTA), and 0.0001 (LDSC).

We further investigate the sensitivity of the screening methods to the choice of screening set size. As an illustrative example, we focus on the HOLP method, which performs best among all screening methods as mentioned above. We vary the screening set size from 10%, 15%, and 20% of the original sample, and the corresponding estimation performance is shown in Figure 5. Following the recommendation of Wang and Leng [2016], we implemented the default rule for determining the number of selected effects, *d*_*s*_ = ⌊*n*_*s*_*/* log *n*_*s*_⌋, where *n*_*s*_ denotes the sample size of the screening set. Both GWASH and LDSC display mild sensitivity to the choice of screening set size, particularly when *κ*_*β*_ is large, whereas GCTA exhibits relatively milder sensitivity.

**Figure 5:**
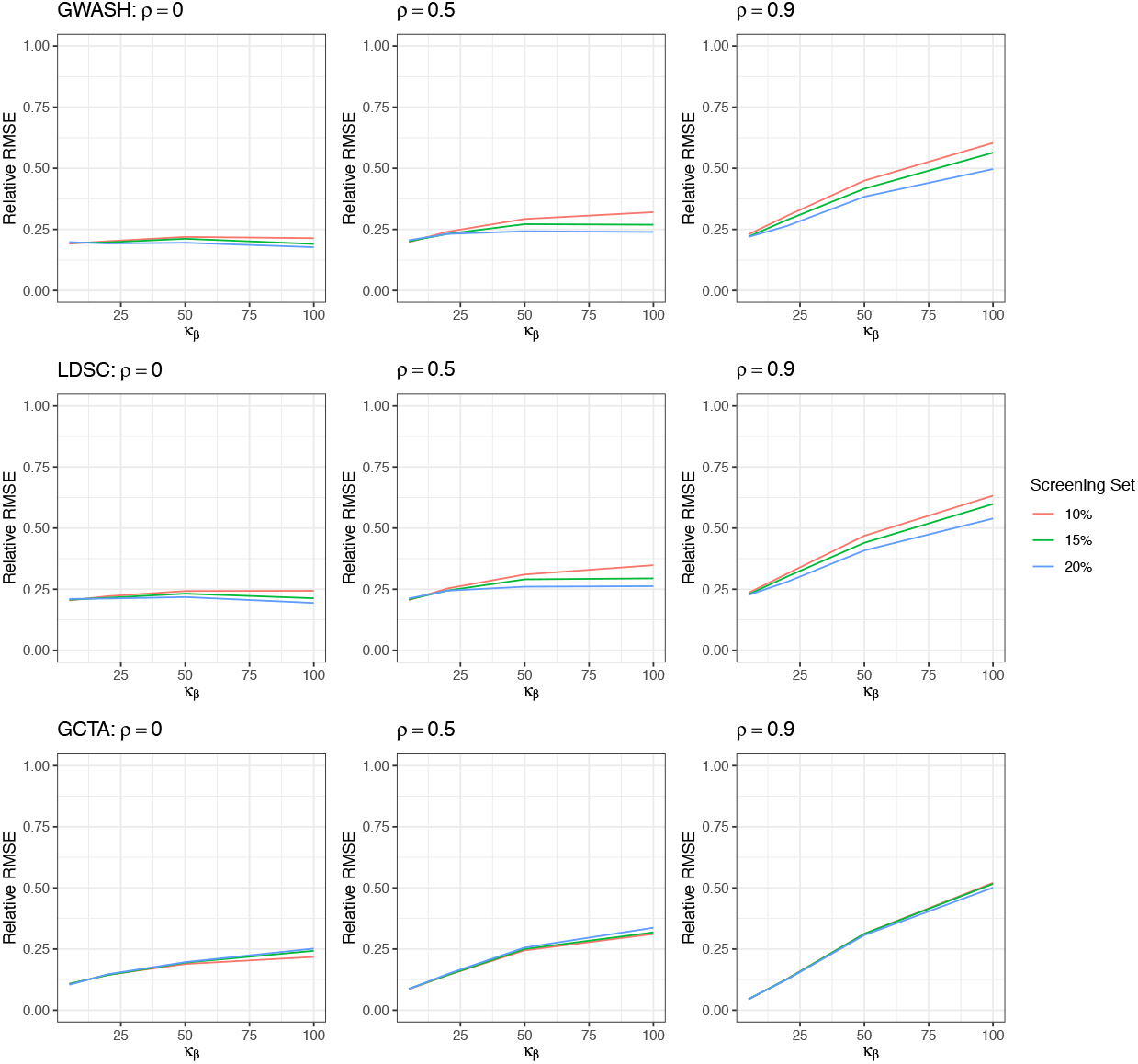
Sensitivity analysis of the HOLP screening procedure using a screening set comprising {10%, 15%, 20%} of the original sample. Results for GWASH, LDSC, GCTA, and LDSC are displayed in the panels from the top to the bottom, respectively. The standard error from the 1000 simulation instances is about 0.001 for the three methods.

## 7 Real Data Analysis

In this section, we apply our proposed FVE decomposition framework to the Adolescent Brain Cognitive Development Study (ABCD; Casey et al. [2018]) to investigate the SNP heritability of our outcome of interest, the PolyVoxel Score (PVS; Loughnan et al. [2022, 2024]), which quantifies the degrees to which an individual’s brain resembles the archetypal hemochromatosis brain. We begin by describing the data preprocessing procedures used in our analysis. Next, we present chromosome-level heritability results and implement a data-driven screening approach to identify strong SNP effect. We then compute the corresponding decomposed FVE estimates using GWASH, LDSC, and GCTA. To evaluate performance, we conducted a bootstrap analysis to determine whether the decomposed estimation provides a statistically significant improvement over the non-decomposed model. Finally, we conduct a sensitivity analysis by varying the size of the screening set used to identify strong effects, using the remaining data for estimation, and comparing the estimation results across different sample sizes.

### 7.1 Data Description and Pre-processing

The ABCD Study provides a comprehensive resource for investigating factors related to child development and mental health. Its genotype data have been widely used in genetic research, including GWAS, with details of the genetic data are available in Fan et al. [2023]. The PolyVoxel Score (PVS) is an imaging-derived measure that quantifies the extent to which an individual’s brain resembles the archetypal hemochromatosis Brain, characterized by regional iron accumulation in motor circuits. Developed using T2-weighted MRI scans, the PVS captures heritable variation associated with brain iron homeostasis. Comprehensive information on the development of the PVS and its integration with the ABCD data is provided in Loughnan et al. [2024].

In our analysis, we included several demographic covariates: age; self-reported ethnicity (categorized as White, Black, Asian, Hispanic, or Other); household income (categorized as “less than $50K,” “$50K–$100K,” and “greater than or equal to $100K”); sex (male or female); proportions of African, European, East Asian, and American genetic ancestry; and the first 10 genetic principal components. The principal components were calculated by Fan et al. [2023] and are included in the analysis following the recommendations of Luo et al. [2021]. In the subsequent analysis, we residualized both the outcome and SNPs on these covariates to remove the population structure, i.e., we regressed the PVS and each SNP on these covariates and used the resulting residuals. A more detailed description of the covariate assessment and processing is provided in Section 2.8.4 of Singh Sachan [2025].

A selection procedure for individuals and SNPs was conducted following Singh Sachan [2025] Section 2.8.3. Due to the longitudinal design of the ABCD Study, we restricted the analysis to baseline observations. For participants with multiple observations, only their baseline data were retained, and individuals missing key variables were excluded. For SNPs, we used imputed genotype data provided by the ABCD Study using the TOPMed imputation server [Taliun et al., 2021], with details described in Fan et al. [2023]. SNPs with a minor allele frequency below 5%, multiallelic variants, those failing Hardy–Weinberg equilibrium, non-autosomal SNPs, and SNPs within the MHC region were excluded. Additional quality control details are provided in Singh Sachan [2025]. The final dataset in our analysis included *n* = 6,623 individuals and *m* = 750,032 SNPs.

### 7.2 Testing the BKE Condition on SNP Effects

We applied the proposed BKE condition test described in Section 5 to each of the 22 chromosomes. To control the family-wise error rate, both the uncorrected *p*-values and a Bonferroni-corrected significance threshold were evaluated. As summarized in Table 1, prior to the multiple testing correction, a notable subset of five chromosomes exhibited significant evidence against the BKE condition. Although these signals did not survive the Bonferroni threshold, the uncorrected results hint at localized deviations from the BKE assumption across multiple chromosomes.

**Table 1:** Original and Bonferroni-adjusted *p*-values of the BKE test across 22 chromosomes (*α* = 0.05).

| CHR | $p$-value | Adj. $p$ | CHR | $p$-value | Adj. $p$ |
| --- | --- | --- | --- | --- | --- |
| 1 | $5.54 \times 10^{-1}$ | 1.00 | 12 | $1.11 \times 10^{-1}$ | 1.00 |
| 2 | $3.84 \times 10^{-1}$ | 1.00 | 13 | $2.14 \times 10^{-1}$ | 1.00 |
| 3 | $9.72 \times 10^{-3}$ | $2.14 \times 10^{-1}$ | 14 | $6.91 \times 10^{-1}$ | 1.00 |
| 4 | $2.59 \times 10^{-2}$ | $5.69 \times 10^{-1}$ | 15 | $3.75 \times 10^{-1}$ | 1.00 |
| 5 | $6.11 \times 10^{-1}$ | 1.00 | 16 | $4.42 \times 10^{-2}$ | $9.72 \times 10^{-1}$ |
| 6 | $7.47 \times 10^{-1}$ | 1.00 | 17 | $1.83 \times 10^{-1}$ | 1.00 |
| 7 | $2.98 \times 10^{-2}$ | $6.56 \times 10^{-1}$ | 18 | $4.85 \times 10^{-1}$ | 1.00 |
| 8 | $1.19 \times 10^{-1}$ | 1.00 | 19 | $3.00 \times 10^{-1}$ | 1.00 |
| 9 | $2.31 \times 10^{-1}$ | 1.00 | 20 | $4.27 \times 10^{-1}$ | 1.00 |
| 10 | $2.99 \times 10^{-1}$ | 1.00 | 21 | $5.32 \times 10^{-2}$ | 1.00 |
| 11 | $8.17 \times 10^{-1}$ | 1.00 | 22 | $6.06 \times 10^{-1}$ | 1.00 |

### 7.3 Decomposed FVE Estimation with Screening Methods

We begin by estimating the FVE using GWASH, LDSC, and GCTA without decomposition, applied to each of the 22 chromosomes separately. The standard error of each GWASH estimate can then be computed using Equation (24) from Schwartzman et al. [2019]. The GCTA software directly computes the corresponding standard errors in its output, while those for LDSC are computed using block jackknife with a block size of 500 SNPs per chromosome. The chromosome-specific FVE estimates obtained using different methods, along with their 95% confidence intervals, are summarized in Figure 6 represented by dashed lines. For most chromosomes, the point estimates obtained from GWASH, LDSC, and GCTA are similar. However, the standard error estimation procedures of GWASH, LDSC, and GCTA differ substantially. In particular, GWASH relies on Dicker’s formula while LDSC employs a block jackknife approach, both of which have been shown to yield inaccurate variance estimates [Pham et al., 2026]. Consequently, the resulting 95% confidence intervals, and hence the statistical significance, can vary across the three methods even on the same chromosome. The intervals reported here, therefore, serve as a heuristic guide rather than a definitive measure of significance.

**Figure 6:**
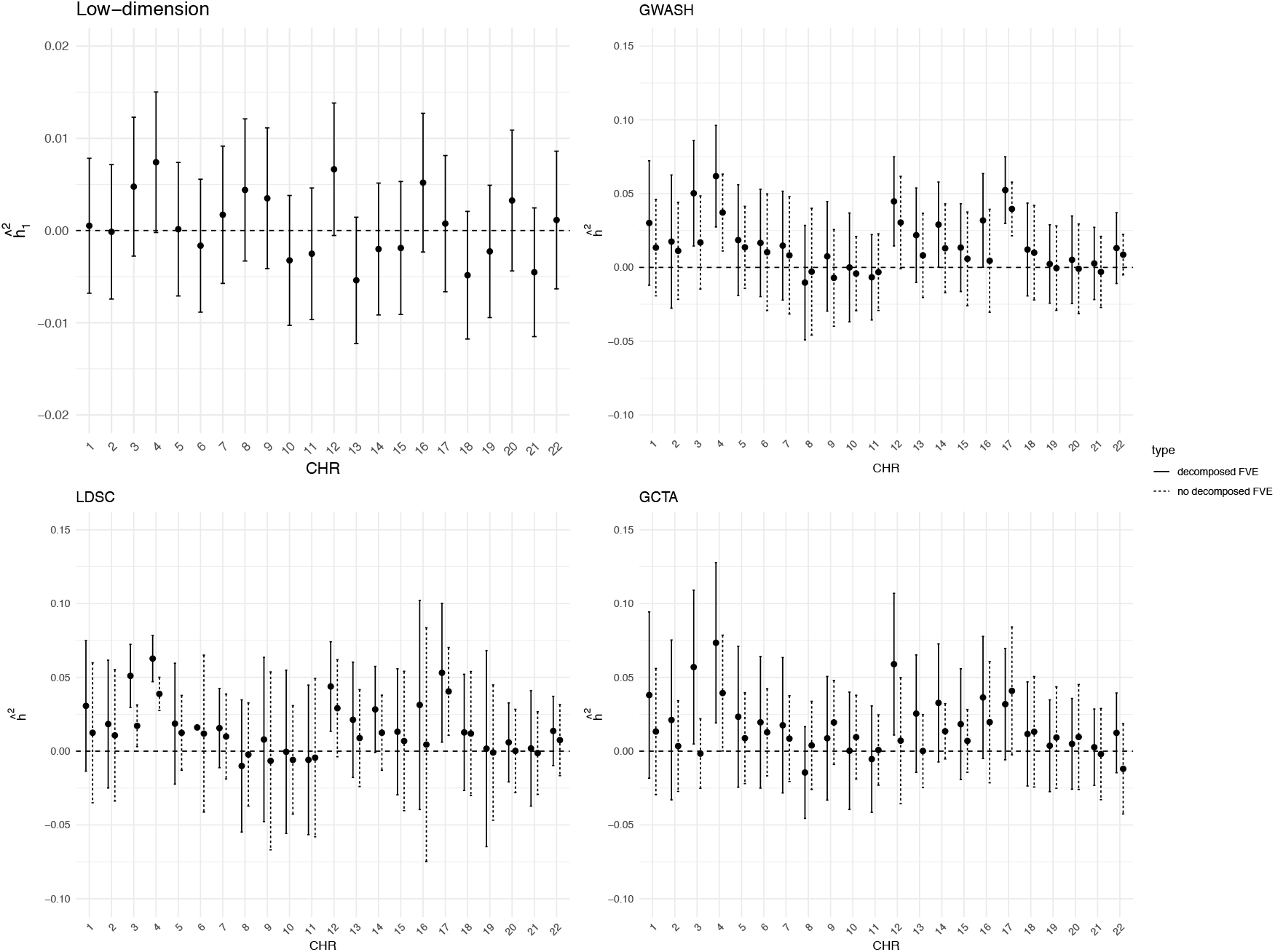
Chromosome-specific FVE estimates based on the decomposed and non-decomposed approaches, where strong effects are identified via the HOLP screening method using 15% of the total sample as the screening set. Confidence intervals are Bonferroni adjusted, controlling the family-wise error rate at *α* = 0.05. Note that the y-axis scales differ across panels.

Loughnan et al. [2024] identified significant SNPs associated with the PVS outcome using a subsample of the UK Biobank, of which 39 SNPs spanning 18 chromosomes were also present in the ABCD dataset. However, this limited number of SNPs is unlikely to capture sufficient strong genetic signal to meaningfully improve the FVE estimation through decomposition. We therefore turn to data-driven screening methods to identify strong-effect SNPs directly from the ABCD dataset.

Specifically, we applied the HOLP method to each chromosome in the ABCD dataset to identify strong SNP effects, as it demonstrated superior performance in the simulation studies. We partitioned the data into a screening set (15%) and an estimation set (85%) to mitigate potential overfitting. HOLP was applied exclusively to the screening set to select SNPs, and decomposed FVE estimates were subsequently obtained using the selected SNPs in the estimation set. Following the recommendation of Wang and Leng [2016], we implemented the default rule for determining the number of selected SNPs, *d*_*s*_ = ⌊*n*_*s*_*/* log *n*_*s*_⌋, where *n*_*s*_ denotes the screening sample size. Under this rule, with 15% of the original sample used as the screening set, we obtained *d*_*s*_ = 143. The estimation results are also presented in Figure 6.

When compared to their non-decomposed counterparts, the decomposed estimates exhibit notable discrepancies at the chromosome level, with the magnitude of these differences varying across both chromosomes and methodologies. Notably, GWASH, LDSC, and GCTA display a high degree of concordance across most chromosomes. Given the similarity between the LDSC and GWASH results, we focus on a more direct comparison between GWASH and GCTA in Figure 7, where the dashed line denotes the identity line (where decomposed estimates equal non-decomposed ones). We also fitted a regression line with fixed intercept at 0 (solid blue line) to compare the decomposed and non-decomposed estimates. For GWASH, the estimated slope of 1.487 (95% CI: [1.224, 1.750]) is significantly greater than 1, while for GCTA the slope of 1.270 (95% CI: [0.700, 1.844]), though wider in uncertainty, is consistent with a similar directional trend. Together, these results point to a systematic pattern in which the decomposed approach yields larger FVE estimates than its non-decomposed counterpart, suggesting that explicitly accounting for strong, localized genetic effects captures additional phenotypic variance that would otherwise be absorbed into or obscured by the polygenic background component. Consequently, the decomposed framework provides a more comprehensive and refined characterization of the genetic architecture of the chromosome level.

**Figure 7:**
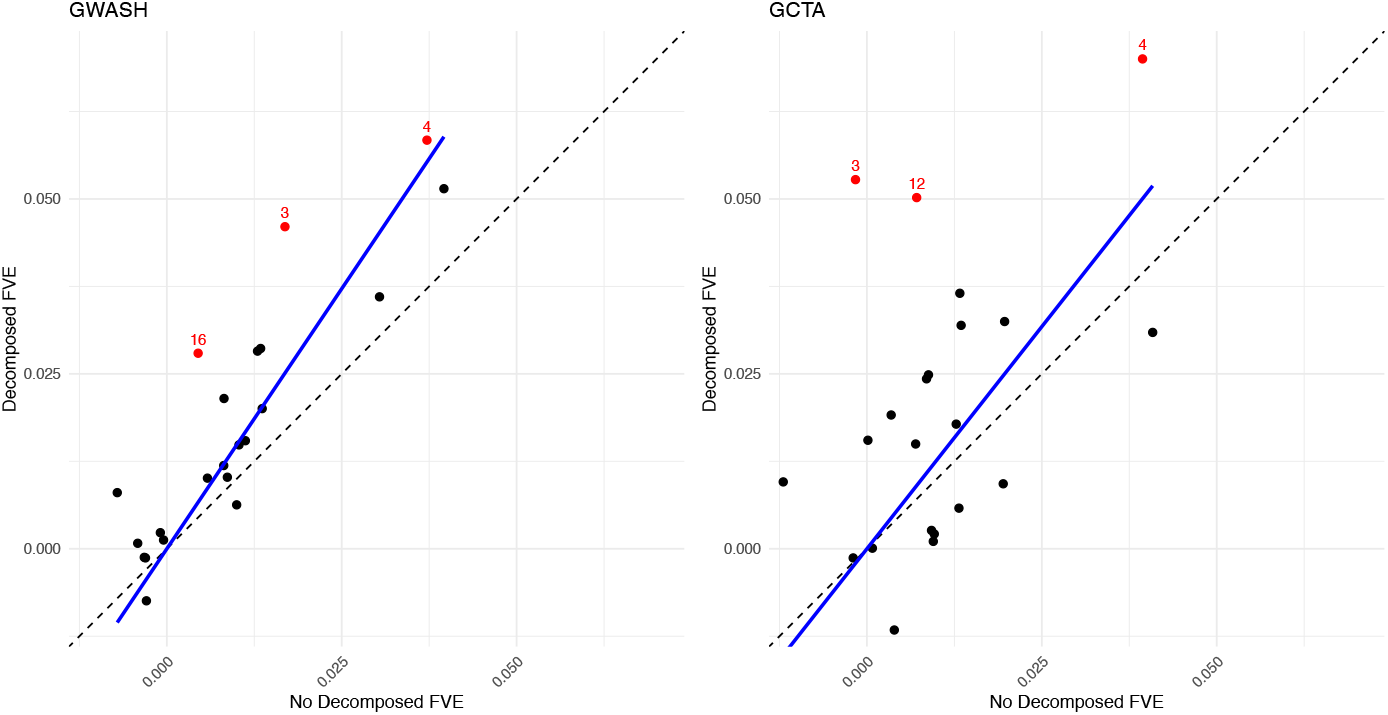
Comparison of decomposed and non-decomposed approaches in GWASH and GCTA for the 22 chromosomes. The dashed line represents the identity line (where decomposed estimates equal non-decomposed estimates), and the blue line is the linear regression fit with a fixed intercept at zero.

The results of the BKE condition tests shown in Table 1 and the estimation results are broadly consistent with each other. Specifically, chromosomes for which the BKE condition is violated (chromosomes 3, 4, 7, and 16) largely overlap with those exhibiting notably positive differences between the decomposed and non-decomposed estimates. This concordance suggests that when the BKE condition is violated, the decomposition leads to meaningfully different FVE estimates, thereby providing empirical support for the practical relevance of the BKE condition test. This pattern further indicates that the identified strong-effect SNPs contribute meaningfully to the PVS phenotype, and that the proposed FVE decomposition procedure provides an effective means of explicitly accounting for such effects in the overall FVE estimation.

Across all three methods, the genome-wide non-decomposed FVE estimates are broadly consistent with one another (Table 2). The decomposed FVE estimates uniformly exceed their non-decomposed counterparts. Specifically, GWASH and LDSC exhibit comparable magnitudes of increase, rising from 0.205 to 0.242 and from 0.206 to 0.249, respectively, while GCTA shows a similar trend, increasing from 0.196 to 0.235. Although the increase in the estimated heritability falls within the confidence interval, the consistent increase observed across all three methods, as well as at the chromosome level (Figure 7), suggests that it is unlikely to be attributable to random noise. Thus, conventional heritability estimates may understate the proportion of phenotypic variance attributable to genetics when strong-effect loci are present, and the decomposed framework consistently recovers a larger share of this variance regardless of the underlying estimation method.

**Table 2:** Genome-wide FVE estimates with 95% confidence intervals for non-decomposed and decomposed approaches across three methods.

| Method | Non-decomposed<br>Estimate (95% CI) | Decomposed<br>Estimate (95% CI) |
| --- | --- | --- |
| GWASH | 0.205 [0.151, 0.260] | 0.242 [0.077, 0.408] |
| LDSC | 0.206 [0.123, 0.289] | 0.249 [0.116, 0.342] |
| GCTA | 0.196 [0.046, 0.345] | 0.235 [0.058, 0.412] |

### 7.4 Sensitivity Analysis

The choice of screening set size represents a trade-off between effective SNP screening and reliable FVE estimation. While a larger estimation set enhances the precision of FVE estimates, a screening set that is too small may lack sufficient statistical power to identify SNPs with strong effects. To assess this balance, we conducted a sensitivity analysis for HOLP by varying the screening–estimation split across proportions 10%, 15%, 20% for screening and the remaining 90%, 85%, 80% for estimation, respectively.

The results are presented in Figure 8, where the estimates for both the low-dimensional and high-dimensional components are shown separately for each method. As the number of SNPs selected by HOLP depends directly on the screening sample size, the corresponding values of *d*_*s*_ were {101, 143, 184} when the screening set was chosen as {10%, 15%, 20%} of the original sample, respectively. For most chromosomes, the decomposed FVE estimates obtained using GWASH, LDSC, and GCTA are consistent across different screening set sizes, with only minor differences in point estimates that remain within an acceptable range. A few exceptions are observed for specific chromosomes, such as chromosomes 1, 5, and 13. Nevertheless, given the overall stability of the results across the majority of chromosomes, the HOLP screening method demonstrates robustness to the choice of screening set size.

**Figure 8:**
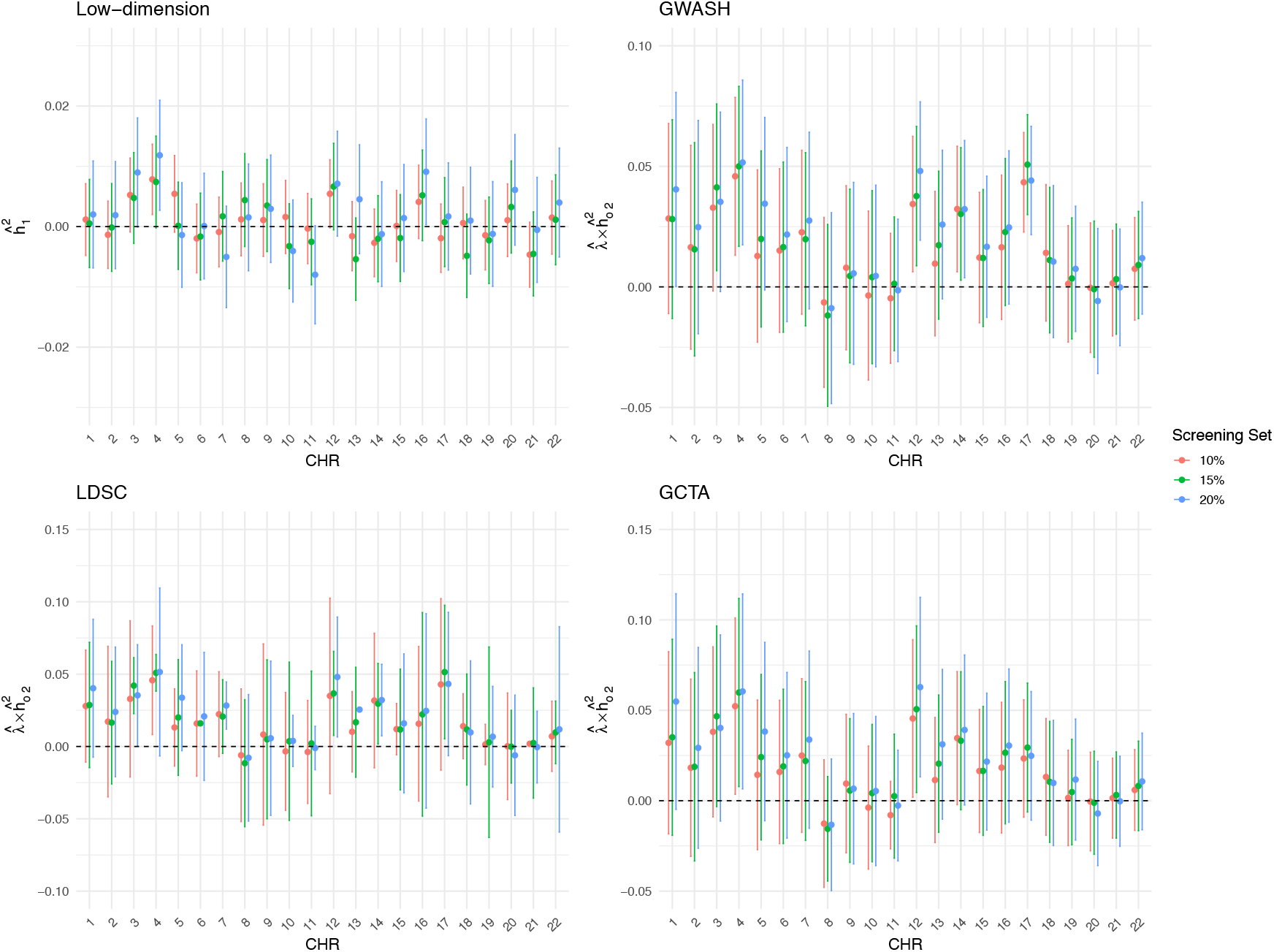
Sensitivity analysis of chromosome-specific decomposed FVE estimates based on strong effects identified by the HOLP screening method using a screening set comprising {10%, 15%, 20%} of the original sample. Note that the y-axis scales differ across the four panels.

## 8 Discussion

We have proposed a novel and general framework for decomposed FVE estimation in high-dimensional linear models where coefficients may have different orders of magnitude. To address limitations of existing methods, such as LDSC, GWASH and GCTA, which rely on the BKE condition, we partition effects into strong and weak components: strong effects, treated as a low-dimensional problem, are estimated via adjusted *R*^2^, while weak effects, after removing and adjusting for the strong ones, are assumed to satisfy the BKE condition and can be estimated using the existing high-dimensional methods. To help illustrate the proposed method and study its performance, we model the coefficients ***β*** using a mixture Gaussian model to control kurtosis and study BKE violations. We further derive a formal relationship between the kurtosis of ***β*** and that of the estimable quantity ***η***, which enables a hypothesis test for evaluating the validity of the BKE assumption. This relationship can also be empirically evaluated under the mixture Gaussian model. Finally, we incorporated several screening procedures into the proposed framework to identify strong effects, thereby facilitating the FVE decomposition. This proposed framework is also successfully applied to the ABCD dataset for chromosome-specific FVE estimation of the polyvoxel score.

Our proposed FVE decomposition framework exhibits superior accuracy in terms of both relative RMSE and bias. In particular, when the correlation among predictors is high and the kurtosis of ***β*** is large, the decomposed estimator substantially reduces both bias and RMSE. The proposed test for the BKE condition demonstrates well-controlled type I error rates and high statistical across different settings. In addition, the incorporated screening procedures effectively identify strong effects, enabling the decomposition to achieve an accurate estimation of the FVE.

There are several limitations and potential directions for future research. First, our estimation framework requires access to the original data and cannot be applied to datasets where only summary statistics are available. Extending the decomposed FVE estimation framework to operate with summary statistics represents a practically relevant avenue for further development. This extension, however, is technically challenging, as summary statistics do not provide sufficient information to perform regression and to residualize the data with respect to the set of SNPs with strong effects.

Second, the standard error estimators for GWASH and LDSC, based on Dicker’s formula and the block jackknife, respectively, are known to provide only approximate variance estimates that may not be fully accurate [Pham et al., 2026]. The confidence intervals reported should therefore be interpreted with care; we leave the development of more accurate procedures for standard error estimation and for inference on the difference between estimators to future work.

## Acknowledgments

The authors thank Chun Chieh Fan and Anubhav Singh-Sachan from UCSD for their help in processing the ABCD data.

## A Further Details on Comparison Methods

### A.1 Genome-wide Complex Trait Analysis (GCTA)

Genome-wide Complex Trait Analysis (GCTA; Yang et al. [2011]) is a widely used framework for estimating the proportion of phenotypic variance explained by genome-wide SNPs, commonly referred to as SNP heritability. Conceptually, GCTA is based on the same underlying linear model as Model (1), which assumes that the observed phenotype can be represented as the sum of additive genetic effects from a large number of SNPs and residual environmental noise.

In the framework of GCTA, all SNP effects are treated as random variables drawn from a normal distribution with a common variance component. This assumption implies that each SNP contributes an equal, infinitesimal amount to the overall genetic variance. The key task is therefore to estimate the variance components corresponding to the genetic and residual sources of variability. GCTA accomplishes this by fitting a linear mixed-effects model using restricted maximum likelihood (REML). Based on these estimates, SNP heritability is then computed as the ratio of the genetic variance to the total phenotypic variance, as defined in Equation (3).

To capture the dependence structure induced by shared genotypes, GCTA constructs a genetic relationship matrix (GRM) from standardized genotype data. The GRM quantifies pairwise genetic similarity between individuals across all SNPs and serves as the covariance matrix for the random genetic effects in the mixed model. Intuitively, the GRM allows GCTA to borrow strength across the entire genome to estimate the aggregate contribution of SNPs, even when individual effects are too small to detect through single-marker association tests.

### A.2 Screening Methods

GWAS are commonly used statistical analyses aimed at identifying genetic variants, typically SNPs, associated with particular traits or diseases. GWAS involves testing each genetic variant individually for association with the phenotype of interest across the genome, often resulting in a massive multiple-testing scenario due to the large number of SNPs examined. Screening methods share conceptual similarities with GWAS in the aim to efficiently identifying significant predictors from large sets of candidates. In our context, screening methods specifically help in isolating predictors with strong effects, which may not be known in advance, facilitating the subsequent application of the decomposed estimation procedure.

Numerous screening methods have been proposed in the literature. Here, we select five representative methods and evaluate their effectiveness. A comparison between these methods can be found in Table 3.

**Table 3:**
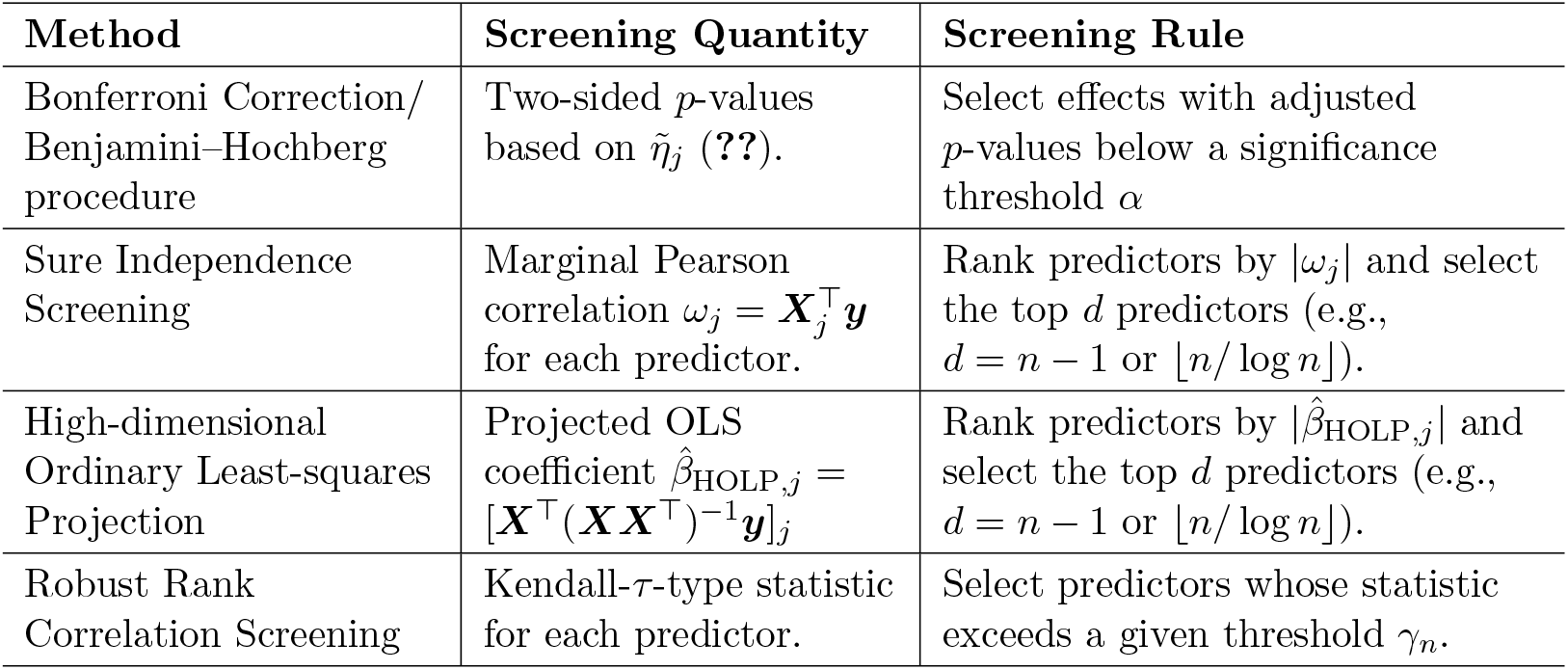
Summary of Selected Screening Methods.

| Method | Screening Quantity | Screening Rule |
| --- | --- | --- |
| Bonferroni Correction/<br>Benjamini–Hochberg<br>procedure | Two-sided $p$-values<br>based on $\tilde{\eta}_j$ (??). | Select effects with adjusted<br>$p$-values below a significance<br>threshold $\alpha$ |
| Sure Independence<br>Screening | Marginal Pearson<br>correlation $\omega_j = \mathbf{X}_j^\top \mathbf{y}$<br>for each predictor. | Rank predictors by $\vert \omega_j\vert $ and select<br>the top $d$ predictors (e.g.,<br>$d = n - 1$ or $\lfloor n/\log n \rfloor$). |
| High-dimensional<br>Ordinary Least-squares<br>Projection | Projected OLS<br>coefficient $\hat{\beta}_{\text{HOLP},j} =$<br>$[\mathbf{X}^\top (\mathbf{X}\mathbf{X}^\top)^{-1} \mathbf{y}]_j$ | Rank predictors by $\vert \hat{\beta}_{\text{HOLP},j}\vert $ and<br>select the top $d$ predictors (e.g.,<br>$d = n - 1$ or $\lfloor n/\log n \rfloor$). |
| Robust Rank<br>Correlation Screening | Kendall-τ-type statistic<br>for each predictor. | Select predictors whose statistic<br>exceeds a given threshold $\gamma_n$. |

- Bonferroni correction [Bonferroni, 1936] or the Benjamini–Hochberg procedure [Benjamini and Hochberg, 1995] are widely used approaches in multiple hypothesis testing and can be applied here to identify potential strong effects. The result in Equation (**??**) states 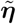 approximately follows a standard normal distribution, 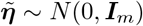. The significance of each parameter 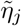 is assessed using a two-sided p-value computed as

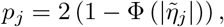

where Φ(·) denotes the cumulative distribution function (CDF) of the standard normal distribution. Since computing *p*_*j*_ for all *j* = 1, …, *m* gives rise to a multiple testing problem, we adjust the p-values {*p*_1_, …, *p*_*m*_} using either the Bonferroni correction or the Benjamini–Hochberg procedure. An effect is deemed strong if its adjusted p-value falls below a pre-specified significance level *α*.
- Sure Independence Screening (SIS, Fan and Lv [2008]) ranks predictors by their marginal Pearson correlations 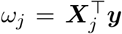, where ***X***_*j*_ is *j*-th column of ***X***, retaining the top *d* (e.g. *d* = ⌊*n/* log *n*⌋ or *n*−1), which provides efficient dimensionality reduction under the assumption that important features exhibit strong marginal effects.
- High-dimensional Ordinary Least-squares Projection (HOLP, Wang and Leng [2016]) computes 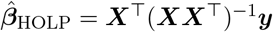. They show that ***X***^⊤^(***XX***^⊤^)^−1^ is the Moore-Penrose inverse of ***X*** when *m > n*; hence this estimator is an extension of the Ordinary Least-squares to the high-dimensional setting where *m > n*. HOLP then ranks variables by 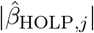 thereby relaxing the “strong marginal correlation” requirement of SIS and improving screening performance when predictors are highly correlated.
- Robust Rank Correlation Screening (RRCS, Li et al. [2012]) replaces Pearson correlation with a Kendall-*τ*–type statistic, as defined in Equation (2.4) of Li et al. [2012], selecting predictors whose statistics exceed a threshold *γ*_*n*_ to enhance robustness against heavy-tailed distributions and nonlinear relationships.

## B Proofs

### B.1 Proof of Proposition 3.1

Based on Equation (14), we have

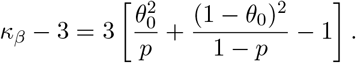

1. Consider scenarios in which the BKE condition is violated:
  - Assume *mp* → *c* and then *m*(1 − *p*) → ∞ as *m* → ∞,

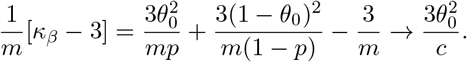
  - Assume *m*(1 − *p*) → *c* and then *mp* → ∞ as *m* → ∞,

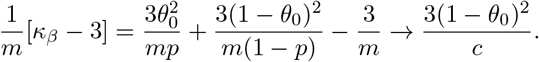
2. Consider scenarios in which the BKE condition is satisfied:
  - *p* = *θ*_0_:

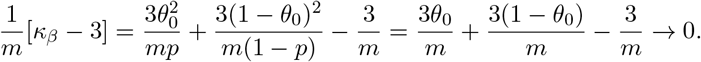
  - *p* ∈ {0, 1} or *θ*_0_ ∈ {0, 1}: In these cases, the mixture Gaussian model (13) collapses to a single normal distribution, yielding *κ*_*β*_ = 3, which trivially satisfies the BKE condition.

### B.2 Proof of Proposition 3.2

Let *κ*_*β*_ = Kurt[*β*_*j*_] and 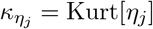. From the definition of *η*_*j*_ in (5),

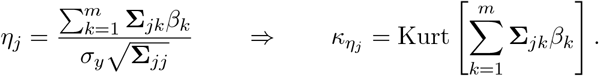

Assuming that the random variables *β*_*k*_ are independent with E[*β*_*k*_] = 0 and 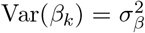, then 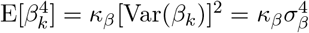. Define

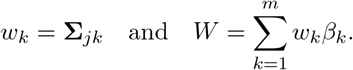

Then

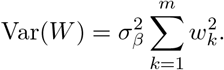

When expanding *W* ^4^, we obtain:

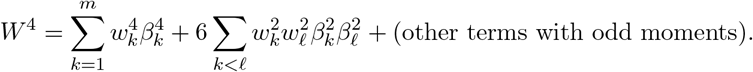

In this expansion of 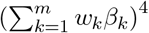, focus on the terms where exactly two distinct indices, say *k* and *ℓ* with *k ≠ ℓ*, appear twice. These terms have the form 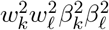. To see the combinatorial factor, note that there are 4 positions in the product. We need to choose 2 of these positions to be occupied by the index *k* (with the remaining two occupied by *ℓ*). The number of ways to choose 2 positions out of 4 is given by the binomial coefficient 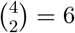. This is the origin of the factor 6 in the expansion.

Taking expectations gives the fourth moment of *W*,

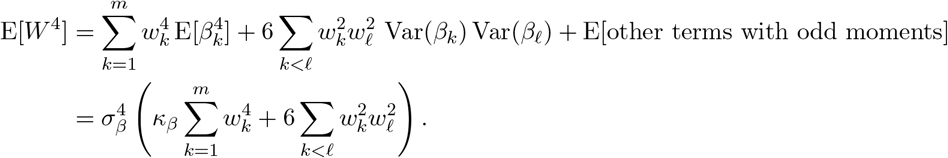

where the terms involving odd powers vanish since E[*β*_*k*_] = 0.

The kurtosis of *W* is

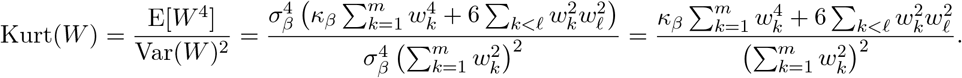

Notice that

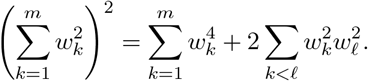

Therefore, the kurtosis can be rewritten as

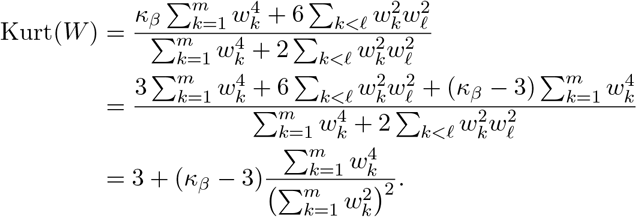

Returning to our original notation with *w*_*k*_ = **Σ**_*jk*_, we have

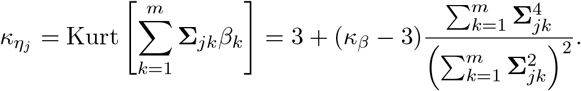

### B.3 Proof of Proposition 4.1

**Step 1:** Recall 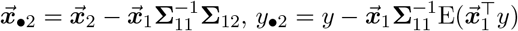, and

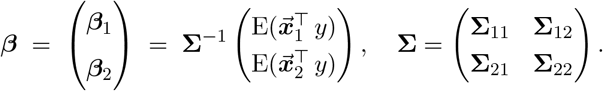

Let

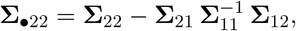

then by the inverse-block formula,

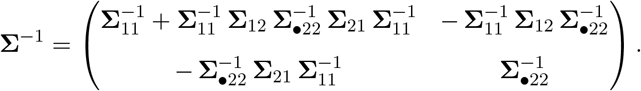

The bottom block ***β***_2_ is given by

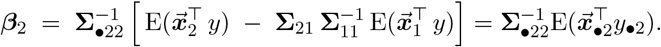

**Step 2:** In order to show Equation (18), it is equivalent to show

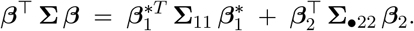

Step 2.1: By definition

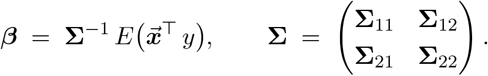

Then we have 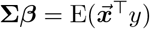, which becomes two vector equations in the block forms:

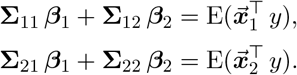

Recall that 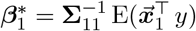. From the first equation above, we can solve for ***β***_1_:

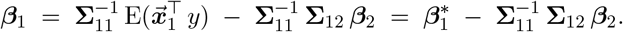

Hence,

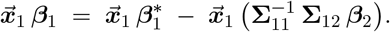

Recall that 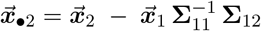. Hence

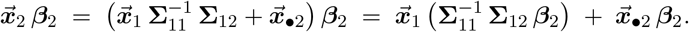

Substituting 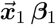 and 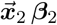 into

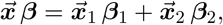

we obtain:

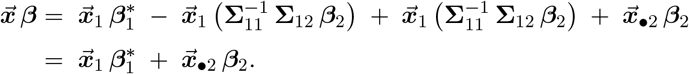

<u>Step 2.2</u>: By definition of 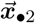, we have

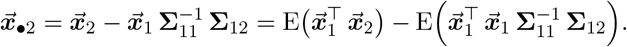

Then we have

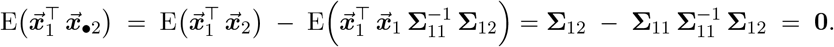

Since 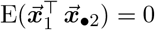,

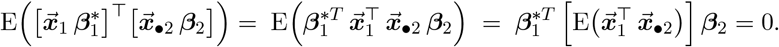

Since 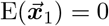 and

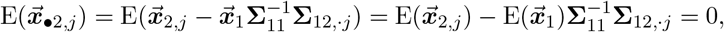

we have

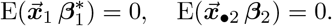

Thus,

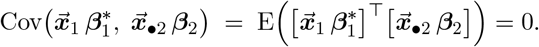

Step 2.3: In

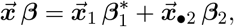

the two terms are uncorrelated. Thus

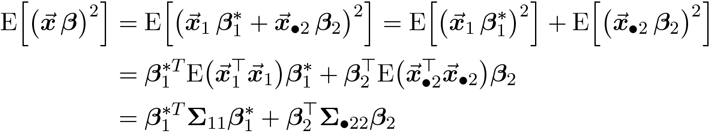

Therefore,

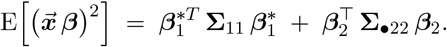

Recall

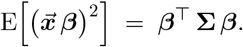

Hence

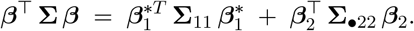

Finally, we have the decomposition of *h*^2^ as

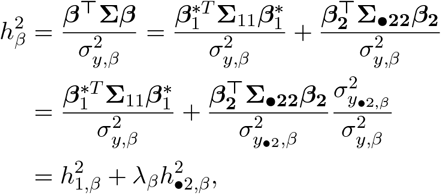

where 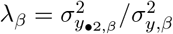.

### B.4 Proof of Proposition 5.1

**Part 1:** Following Model (1), we have E[***y***|***X, β***] = ***Xβ*** and 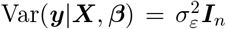. With the sample covariance matrix of ***X*** is 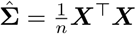 and the correlation 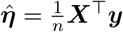:

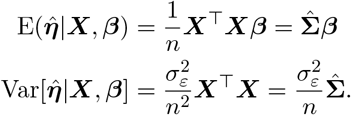

With the assumption that E(***β***) = 0 and Var[***β***] = *h*^2^***I****/m*, by using the law of total variance, we can have the expression of the covariance matrix ***V*** given ***X***:

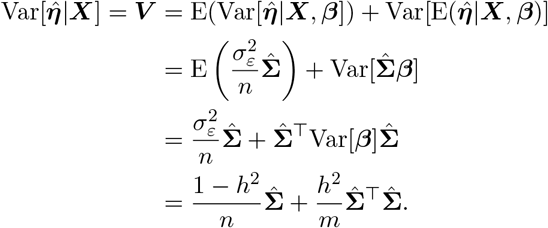

**Part 2:** Substituting the model ***y*** = ***Xβ*** + ***ε***, we have

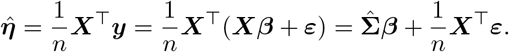

Since we assume 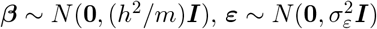 and 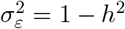, with ***β*** being independent of ***ε*** in Model (1), we have

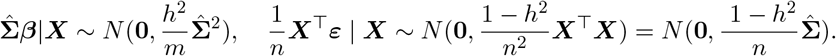

As both terms are Gaussian and independent given ***X***, it follows that their summation 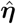 is also Gaussian conditional on ***X***:

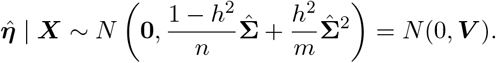

## C Additional Simulation Results

The estimation results of LDSC are very similar to those of GWASH; the estimation performance of LDSC in terms of relative RMSE is shown in Figure 10. The estimation results of the decomposed GCTA estimator, in comparison to the original GCTA estimator, are presented in Figure 9 for relative RMSE. The performance of the GCTA-based decomposition closely mirrors that of the GWASH-based decomposition. In high-dimensional settings, the decomposed estimator does not exhibit a clear advantage over the original GCTA estimator when the kurtosis of ***β*** is relatively low (*κ*_*β*_ = 5). However, as the kurtosis increases (*κ*_*β*_ ≥ 20), the decomposed estimator achieves a substantially lower RMSE. These findings suggest that GCTA is sensitive to heavy-tailed distributions and that our proposed decomposition framework can help mitigate this issue.

**Figure 9:**
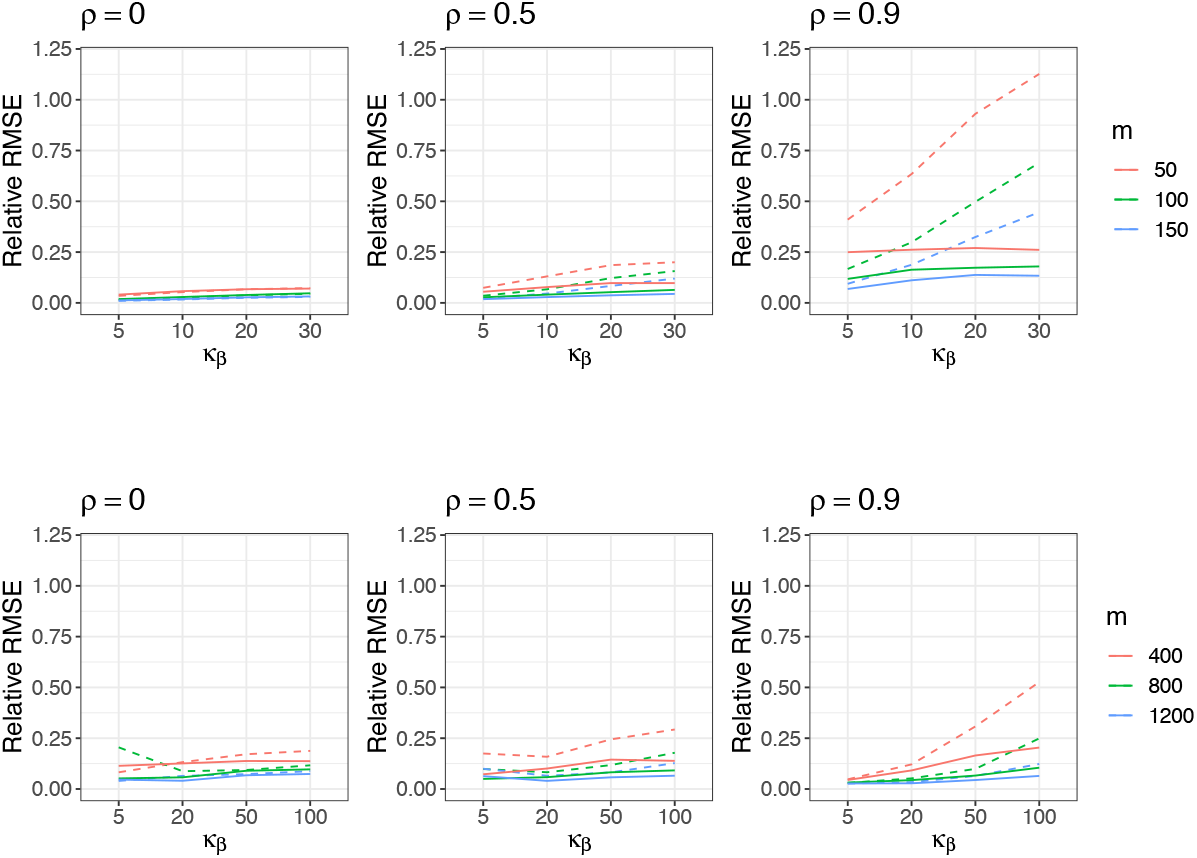
Estimation performance of the GCTA without decomposition (dashed line) and decomposed GCTA (solid line) under the high-dimension setting. The standard error from 1000 simulation instances is about 0.00015 (low-dimension) and 0.0002 (high-dimension).

**Figure 10:**
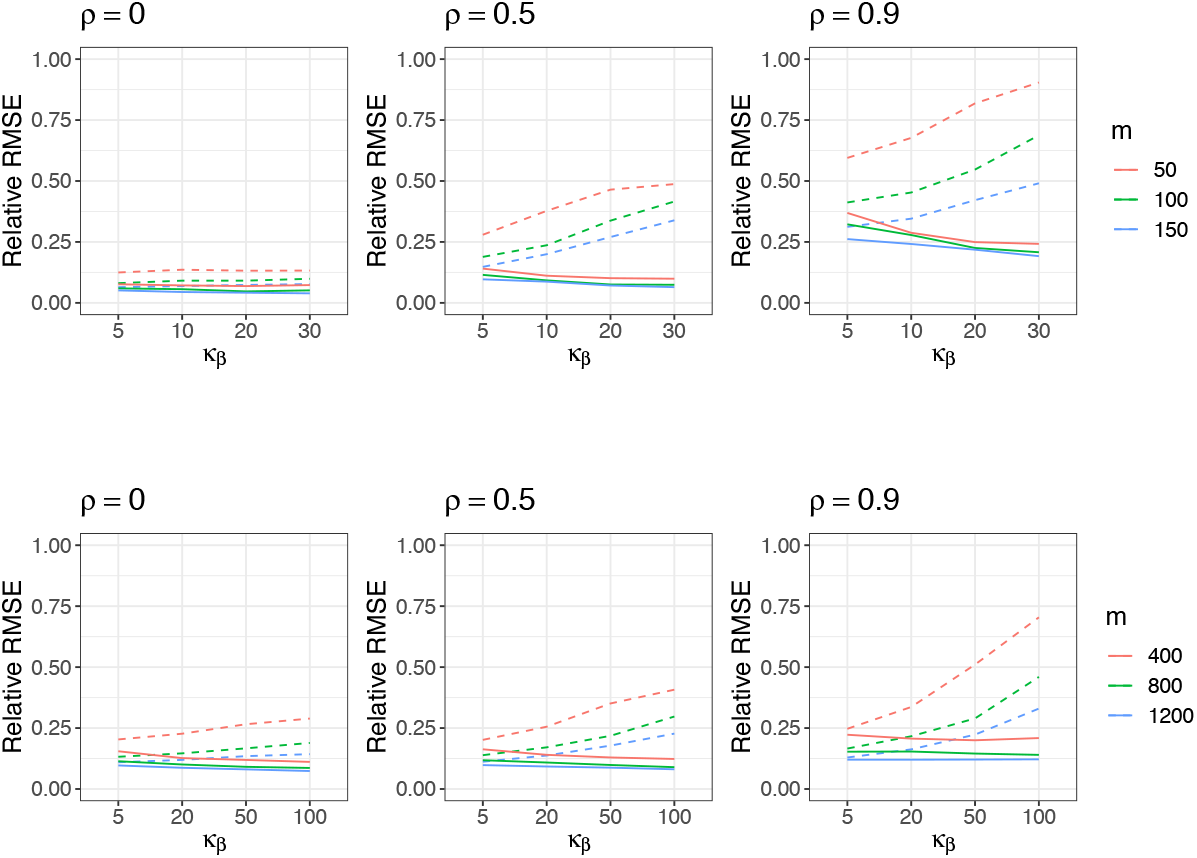
Estimation performance of the LDSC without decomposition (dashed line) and decomposed LDSC (solid line) under the high-dimension setting. The standard error from 1000 simulation instances is about 0.0003 (low-dimension) and 0.0006 (high-dimension).

As shown in Figure 11, 12, 13, in both low- and high-dimensional settings, the decomposed estimates with GWASH, LDSC, and GCTA consistently exhibit smaller bias than the non-decomposed estimates.

**Figure 11:**
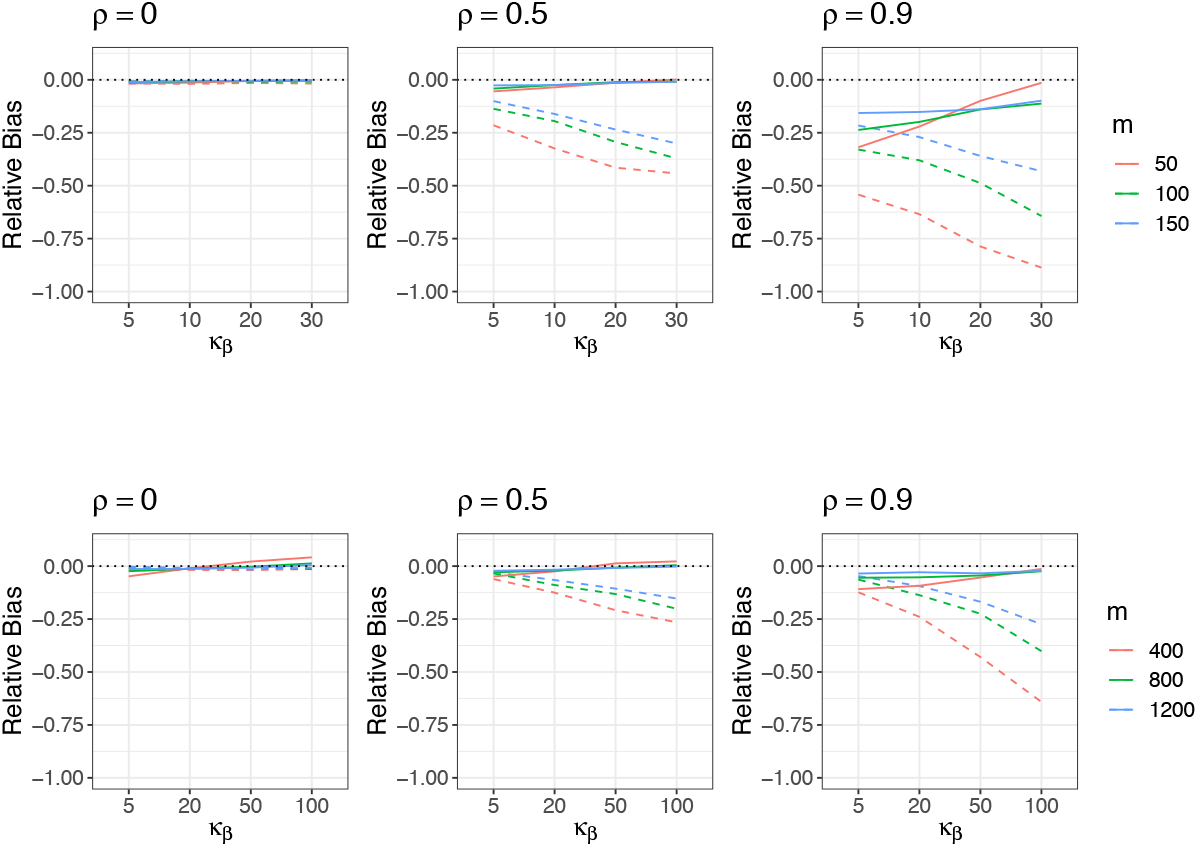
Estimation performance of GWASH without decomposition (dashed line) and decomposed GWASH (solid line) in the low-dimensional setting (top row) and high-dimensional setting (bottom row). The standard error from 1000 simulation instances is about 0.002 (low-dimension) and 0.003 (high-dimension).

**Figure 12:**
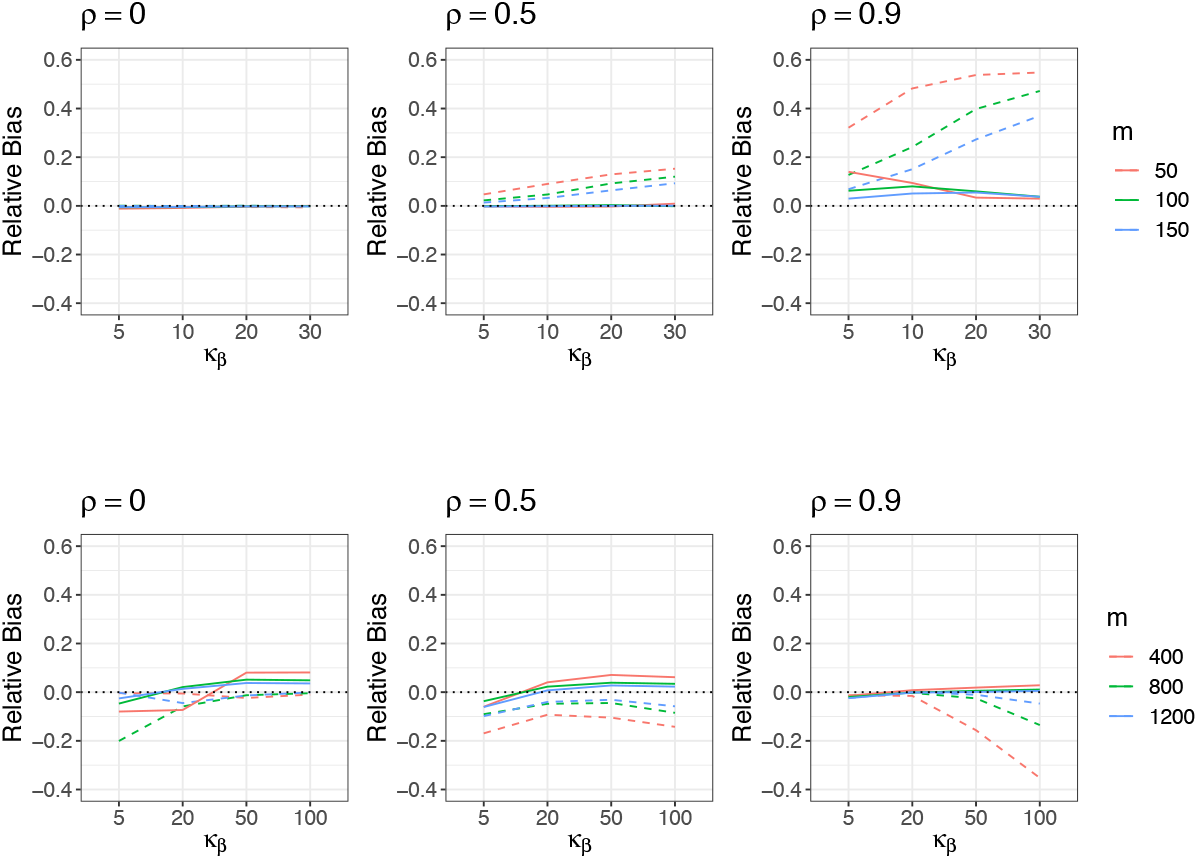
Estimation performance of the GCTA without decomposition (dashed line) and decomposed GCTA (solid line) under the high-dimension setting. The standard error from 1000 simulation instances is about 0.0015 (low-dimension) and 0.0018 (high-dimension).

**Figure 13:**
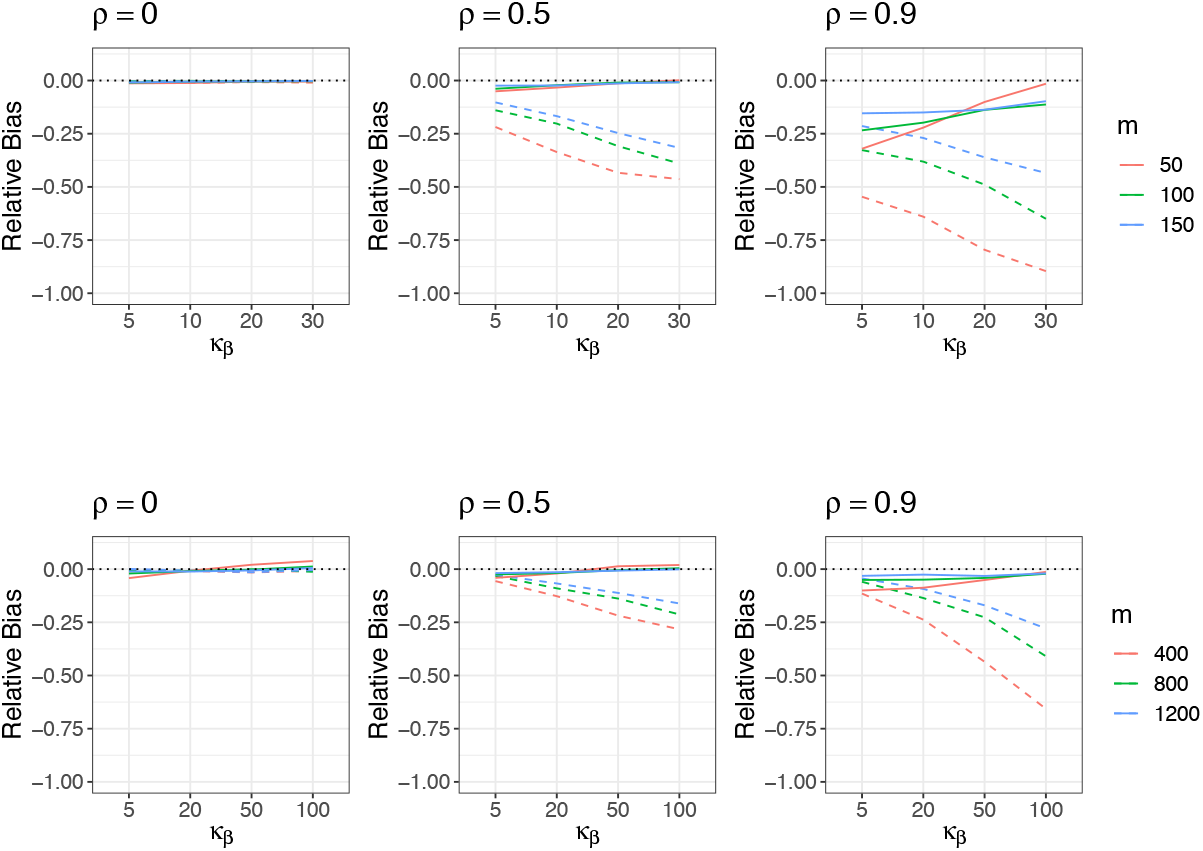
Estimation performance of the LDSC without decomposition (dashed line) and decomposed LDSC (solid line) under the high-dimension setting. The standard error from 1000 simulation instances is about 0.002 (low-dimension) and 0.004 (high-dimension).

## References

David Azriel, Samuel Davenport, and Armin Schwartzman. Consistency of heritability estimation from summary statistics in high-dimensional linear models. arXiv preprint 2502.11144, 2025.

Yoav Benjamini and Yosef Hochberg. Controlling the false discovery rate: a practical and powerful approach to multiple testing. Journal of the Royal statistical society: series B (Methodological), 57(1):289–300, 1995.

Carlo Bonferroni. Teoria statistica delle classi e calcolo delle probabilita. Pubblicazioni del R istituto superiore di scienze economiche e commericiali di firenze, 8:3–62, 1936.

Brendan K Bulik-Sullivan, Po-Ru Loh, Hilary K Finucane, Stephan Ripke, Jian Yang, Schizophrenia Working Group of the Psychiatric Genomics Consortium, Nick Patterson, Mark J Daly, Alkes L Price, and Benjamin M Neale. LD score regression distinguishes confounding from polygenicity in genome-wide association studies. Nature genetics, 47(3):291–295, 2015.

Betty Jo Casey, Tariq Cannonier, May I Conley, Alexandra O Cohen, Deanna M Barch, Mary M Heitzeg, Mary E Soules, Theresa Teslovich, Danielle V Dellarco, Hugh Garavan, et al. The adolescent brain cognitive development (ABCD) study: imaging acquisition across 21 sites. Developmental cognitive neuroscience, 32:43–54, 2018.

Lee H Dicker. Variance estimation in high-dimensional linear models. Biometrika, 101(2):269–284, 2014.

Luke M Evans, Rasool Tahmasbi, Scott I Vrieze, Gonçalo R Abecasis, Sayantan Das, Steven Gazal, Douglas W Bjelland, Teresa R De Candia, Haplotype Reference Consortium, Michael E Goddard, et al. Comparison of methods that use whole genome data to estimate the heritability and genetic architecture of complex traits. Nature genetics, 50(5):737–745, 2018.

Chun Chieh Fan, Robert Loughnan, Sylia Wilson, John K Hewitt, and ABCD Genetic Working Group. Genotype data and derived genetic instruments of adolescent brain cognitive development study^®^ for better understanding of human brain development. Behavior genetics, 53(3):159–168, 2023.

Jianqing Fan and Jinchi Lv. Sure independence screening for ultrahigh dimensional feature space. Journal of the Royal Statistical Society Series B: Statistical Methodology, 70(5):849–911, 2008.

Hilary K Finucane, Brendan Bulik-Sullivan, Alexander Gusev, Gosia Trynka, Yakir Reshef, Po-Ru Loh, Verneri Anttila, Han Xu, Chongzhi Zang, Kyle Farh, et al. Partitioning heritability by functional annotation using genome-wide association summary statistics. Nature genetics, 47(11): 1228–1235, 2015.

Ronald A Fisher. XV.—The correlation between relatives on the supposition of mendelian inheritance. Earth and Environmental Science Transactions of the Royal Society of Edinburgh, 52(2):399–433, 1919.

Alexander Gusev, S Hong Lee, Gosia Trynka, Hilary Finucane, Bjarni J Vilhjálmsson, Han Xu, Chongzhi Zang, Stephan Ripke, Brendan Bulik-Sullivan, Eli Stahl, et al. Partitioning heritability of regulatory and cell-type-specific variants across 11 common diseases. The American Journal of Human Genetics, 95(5):535–552, 2014.

Dominic Holland, Oleksandr Frei, Rahul Desikan, Chun-Chieh Fan, Alexey A Shadrin, Olav B Smeland, Vijay S Sundar, Paul Thompson, Ole A Andreassen, and Anders M Dale. Beyond snp heritability: Polygenicity and discoverability of phenotypes estimated with a univariate gaussian mixture model. PLoS Genetics, 16(5):e1008612, 2020.

Kangcheng Hou, Kathryn S Burch, Arunabha Majumdar, Huwenbo Shi, Nicholas Mancuso, Yue Wu, Sriram Sankararaman, and Bogdan Pasaniuc. Accurate estimation of SNP-heritability from biobank-scale data irrespective of genetic architecture. Nature genetics, 51(8):1244–1251, 2019.

Lucas Janson, Rina Foygel Barber, and Emmanuel Candes. Eigenprism: inference for high dimensional signal-to-noise ratios. Journal of the Royal Statistical Society Series B: Statistical Methodology, 79(4):1037–1065, 2017.

Vincent Laville, Jae H Kang, Clara C Cousins, Adriana I Iglesias, Réka Nagy, Jessica N Cooke Bailey, Robert P Igo Jr, Yeunjoo E Song, Daniel I Chasman, William G Christen, et al. Genetic correlations between diabetes and glaucoma: an analysis of continuous and dichotomous phenotypes. American journal of ophthalmology, 206:245–255, 2019.

Sang Hong Lee, Naomi R Wray, Michael E Goddard, and Peter M Visscher. Estimating missing heritability for disease from genome-wide association studies. The American Journal of Human Genetics, 88(3):294–305, 2011.

Gaorong Li, Heng Peng, Jun Zhang, and Lixing Zhu. Robust rank correlation based screening. The Annals of Statistics, 40(3):1846–1877, 2012.

Robert Loughnan, Jonathan Ahern, Mary Boyle, Terry L Jernigan, Donald J Hagler Jr, John R Iversen, Oleksandr Frei, Diana M Smith, Ole Andreassen, Noah Zaitlen, et al. Hemochromatosis neural archetype reveals iron disruption in motor circuits. Science Advances, 10(47):eadp4431, 2024.

Robert J Loughnan, Alexey A Shadrin, Oleksandr Frei, Dennis van der Meer, Weiqi Zhao, Clare E Palmer, Wesley K Thompson, Carolina Makowski, Terry L Jernigan, Ole A Andreassen, et al. Generalization of cortical mostest genome-wide associations within and across samples. Neuroimage, 263:119632, 2022.

Monica Luciana, James M Bjork, Bonnie J Nagel, Deanna M Barch, Raul Gonzalez, Sara Jo Nixon, and Marie T Banich. Adolescent neurocognitive development and impacts of substance use: Overview of the adolescent brain cognitive development (ABCD) baseline neurocognition battery. Developmental cognitive neuroscience, 32:67–79, 2018.

Yang Luo, Xinyi Li, Xin Wang, Steven Gazal, Josep Maria Mercader, Me Research Team, SIGMA Type 2 Diabetes Consortium, Benjamin M Neale, Jose C Florez, Adam Auton, et al. Estimating heritability and its enrichment in tissue-specific gene sets in admixed populations. Human molecular genetics, 30(16):1521–1534, 2021.

Michael Lynch, Bruce Walsh, et al. Genetics and analysis of quantitative traits, volume 1. Sinauer Sunderland, MA, 1998.

Benjamin K Pham, Samuel Davenport, David Azriel, and Armin Schwartzman. When can whole-genome snp heritability be reliably estimated from summary statistics? bioRxiv, pages 2026–05, 2026.

Tinca JC Polderman, Beben Benyamin, Christiaan A De Leeuw, Patrick F Sullivan, Arjen Van Bo- choven, Peter M Visscher, and Danielle Posthuma. Meta-analysis of the heritability of human traits based on fifty years of twin studies. Nature genetics, 47(7):702–709, 2015.

Armin Schwartzman, Andrew J Schork, Rong Zablocki, and Wesley K Thompson. A simple, consistent estimator of snp heritability from genome-wide association studies. The annals of applied statistics, 13(4):2509, 2019.

Huwenbo Shi, Gleb Kichaev, and Bogdan Pasaniuc. Contrasting the genetic architecture of 30 complex traits from summary association data. The American Journal of Human Genetics, 99(1): 139–153, 2016.

Anubhav Nikunj Singh Sachan. Estimating genetic effects under different data complexities. PhD thesis, UC San Diego, 2025.

Doug Speed and David J Balding. Sumher better estimates the snp heritability of complex traits from summary statistics. Nature genetics, 51(2):277–284, 2019.

Doug Speed, Gibran Hemani, Michael R Johnson, and David J Balding. Improved heritability estimation from genome-wide snps. The American Journal of Human Genetics, 91(6):1011–1021, 2012.

Patrick F Sullivan and Daniel H Geschwind. Defining the genetic, genomic, cellular, and diagnostic architectures of psychiatric disorders. Cell, 177(1):162–183, 2019.

Daniel Taliun, Daniel N Harris, Michael D Kessler, Jedidiah Carlson, Zachary A Szpiech, Raul Torres, Sarah A Gagliano Taliun, André Corvelo Stephanie M Gogarten, Hyun Min Kang, et al. Sequencing of 53,831 diverse genomes from the NHLBI TOPMed Program. Nature, 590(7845): 290–299, 2021.

Peter M Visscher, Naomi R Wray, Qian Zhang, Pamela Sklar, Mark I McCarthy, Matthew A Brown, and Jian Yang. 10 years of gwas discovery: biology, function, and translation. The American Journal of Human Genetics, 101(1):5–22, 2017.

Xiangyu Wang and Chenlei Leng. High dimensional ordinary least squares projection for screening variables. Journal of the Royal Statistical Society Series B: Statistical Methodology, 78(3):589–611, 2016.

Jichun Xie, T Tony Cai, John Maris, and Hongzhe Li. False discovery rate control for high-dimensional dependent data with an application to large-scale genetic association studies. The Annals of Applied Statistics, 5(2A):1207–1225, 2011.

Jian Yang, Beben Benyamin, Brian P McEvoy, Scott Gordon, Anjali K Henders, Dale R Nyholt, Pamela A Madden, Andrew C Heath, Nicholas G Martin, Grant W Montgomery, et al. Common SNPs explain a large proportion of the heritability for human height. Nature genetics, 42(7): 565–569, 2010.

Jian Yang, S Hong Lee, Michael E Goddard, and Peter M Visscher. GCTA: a tool for genome-wide complex trait analysis. The American Journal of Human Genetics, 88(1):76–82, 2011.

